# The cellular characterisation of SARS-CoV-2 spike protein in virus-infected cells using Receptor Binding Domain-binding specific human monoclonal antibodies

**DOI:** 10.1101/2021.12.06.471528

**Authors:** Conrad En-Zuo Chan, Ching-Ging Ng, Angeline Pei-Chew Lim, Shirley Lay-Kheng Seah, De-Hoe Chye, Steven Ka-Khuen Wong, Jie-Hui Lim, Vanessa Zi-Yun Lim, Soak-Kuan Lai, Pui-San Wong, Kok-Mun Leong, Yi-Chun Liu, Richard J Sugrue, Boon-Huan Tan

**Affiliations:** Biological Defence Programme, DSO National Laboratories, 27 Medical Drive, Singapore 117510, Singapore; Infectious Disease Laboratory, National Centre for Infectious Diseases, 16 Jalan Tan Tock Seng, Singapore 308442; School of Biological Sciences, 60 Nanyang Drive, Nanyang Technological University, Singapore 637551, Singapore; Infection and Immunity, LKC School of Medicine, Nanyang Technological University, 11 Mandalay Road, Singapore 308232, Singapore

**Keywords:** SARS-CoV-2, S protein, receptor binding domain (RBD), Human monoclonal antibodies, convalescent patient serum, furin, cell-to-cell transmission

## Abstract

A human monoclonal antibody panel (PD4, PD5, PD7, SC23 and SC29) was isolated from the B cells of convalescent patients and used to examine the S protein in SARS-CoV-2- infected cells. While all five antibodies bound conformational-specific epitopes within SARS-CoV-2 Spike (S) protein, only PD5, PD7, and SC23 were able to bind to the Receptor Binding Domain (RBD). Immunofluorescence microscopy was used to examine the S protein RBD in cells infected with the Singapore isolates SARS-CoV-2/0334 and SARS-CoV-2/1302. The RBD-binders exhibited a distinct cytoplasmic staining pattern that was primarily localised within the Golgi complex and was distinct from the diffuse cytoplasmic staining pattern exhibited by the non-RBD binders (PD4 and SC29). These data indicated that the S protein adopted a conformation in the Golgi complex that enabled the RBD recognition by the RBD-binders. The RBD-binders also recognised the uncleaved S protein indicating that S protein cleavage was not required for RBD recognition. Electron microscopy indicated high levels of cell-associated virus particles, and multiple cycle virus infection using RBD-binder staining provided evidence for direct cell-to-cell transmission for both isolates. Although similar levels of RBD-binder staining was demonstrated for each isolate, the SARS-CoV-2/1302 exhibited slower rates of cell-to-cell transmission. These data suggest that a conformational change in the S protein occurs during its transit through the Golgi complex that enables RBD recognition by the RBD-binders, and suggests that these antibodies can be used to monitor S protein RBD formation during the early stages of infection.

**Importance:** The SARS CoV-2 spike (S) protein receptor binding domain (RBD) mediates the attachment of SARS CoV-2 to the host cell. This interaction plays an essential role in initiating virus infection and the S protein RBD is therefore a focus of therapeutic and vaccine interventions. However, new virus variants have emerged with altered biological properties in the RBD that can potentially negate these interventions. Therefore an improved understanding of the biological properties of the RBD in virus-infected cells may offer future therapeutic strategies to mitigate SARS CoV-2 infection. We used physiologically relevant antibodies that were isolated from the B cells of convalescent COVID19 patients to monitor the RBD in cells infected with SARS CoV-2 clinical isolates. These immunological reagents specifically recognise the correctly folded RBD and were used to monitor the appearance of the RBD in SARS CoV-2-infected cells and identified the site where the RDB first appears.

## Introduction

Coronavirus disease 2019 (COVID-19) is a new respiratory-borne infectious disease caused by severe acute respiratory syndrome coronavirus 2 (SARS-CoV-2) [1, 2]. Since its first formal identification in Wuhan City, Hubei Province in China in 2019, SARS-CoV-2 has been responsible for approximately 255 million infections and 5.1 million deaths worldwide (https://covid19.who.int/ assessed on 19^th^ November 2021). The SARS-CoV-2 virus belongs to the *Coronaviridae* family, which includes both established coronaviruses that usually cause mild to moderate respiratory disease in human (e.g. the human coronavirus 229E) and newer emerging viruses that are associated with high mortality rates in humans (e.g. severe acute respiratory syndrome coronavirus 1) [3–5]. The SARS-CoV-2 like other coronaviruses contains a positive-sense, non-segmented and single-stranded RNA genome (vRNA) and contains between ten to fourteen open reading frames (ORF), which encode various virus proteins in the order: 5’UTR, replicase (ORF1a/1b), spike (S), envelope (E), membrane (M), nucleocapsid (N), 3’UTR and poly A tail [6]. The mature SARS-CoV-2 particle is surrounded by a lipid envelope that is derived from the host cell, and into which the M, E and S proteins are inserted. Since the first published sequence of the SARS-CoV-2 that was isolated in Wuhan (WIV04) [7], new sequence variants of SARS-CoV-2 have been identified (e.g. alpha, beta, gamma and delta variants) that exhibit variations in the virus genome sequences that change the biological properties of the virus (https://www.who.int/en/activities/tracking-SARS-CoV-2-variants/ accessed on 20^th^ November 2021). It is postulated that these changes may also lead to increased virus transmission and lead to altered immunogenicity in humans [8–10]. Therefore, an improved understanding of the biology of new and existing circulating SARS-CoV-2 variants is required to better understand the risks that these variants pose.

The S protein mediates the entry of the virus into the host cell and it thus plays an essential role in initiating the virus infection. The S protein exists as a homotrimer and protrudes from the virus envelope as an array of club-like projections. It is a dominant feature on the surface of virus particle, and this topology is a defining feature in coronavirus identification using diagnostic electron microscopy. The virus envelope surrounds internal virus structures, such as the virus nucleocapsid that is formed by the association between the vRNA and the nucleocapsid (N) protein. The entry of the SARS-CoV-2 genome into susceptible cells occurs by fusion of the virus envelope and cell membrane, and the S protein mediates both the cell attachment (via the cell receptor), and the fusion of the virus and cell membranes. The S protein is initially synthesised as a single polypeptide chain (S0), and it is subsequently cleaved into the S1 and S2 subunits at two cleavage sites. The S1 domain contains the receptor binding domain (RBD) that mediates binding of the virus to the target cell via angiotensin-converting enzyme 2 (ACE2) host cell receptor. The S2 domain is anchored to the virus envelope by a transmembrane domain, and it contains the fusion peptide and heptad repeat regions that mediate the process of membrane fusion. Unlike the SARS-CoV-1 S protein, the cleavage site of the S protein of the SARS-CoV-2 contains the sequence RRAR at the S1/S2 site which is recognized by the ubiquitous cellular protease furin (reviewed in [11]). A second cleavage site within the S2 domain (the S2ˈsite) is required during the early stages of virus cell entry, however cleavage at the S1/S2 site is required for S2ˈcleavage [12]. Although cleavage at the S2ˈsite is mediated by transmembrane proteases serine 2 (TMPRSS2), TMPRSS2 is not expressed in Vero E6 cells [13] and it is proposed that the S2ˈcleavage site can also be processed by other cellular proteases in these cells [14].

The S protein has also become a focus in the development of antiviral drug strategies using small molecule inhibitors and passive immunisation using human monoclonal antibodies (hMAbs) [15]. Given the importance of the S protein RBD during virus cell attachment, these immunological reagents often target the RBD to prevent the initial stages of SARS-CoV-2 entry into host cells. The isolation of hMAbs from the B cells of convalescent serum can be used in passive immunization for the timely treatment and prevention of SARS-CoV-2 infection, and the discovery of hMAbs from COVID19 convalescent patients have shown therapeutic potential [16–19]. A number of these have been granted emergency authorisation and progressed through clinical trials for use as antibodies therapeutics [20, 21]. Although the emphasis has been on the development of reagents that can block the RBD, the potential use of non-RBD binders with virus neutralising activity as part of a cocktail of antibodies therapeutics is also expected to overcome problems associated with the emergence of new virus variants [17]. The most important practical criterion for these immunological reagents is that they neutralize virus infection by e.g. blocking attachment of the virus to the host cell. However, it is also expected that these reagents could also provide other useful information about the biology of the S protein of existing and newly emerged SARS-CoV-2 variants. The individual monoclonal antibodies could also potentially form part of a wider polyclonal immune response during SARS-CoV-2 infection, and characterizing the individual monoclonal antibodies could provide useful information about how these antibody responses interact with the virus. We have previously described a panel of S protein hMAbs that were isolated using the B cells of convalescent COVID 19 patients in Singapore [19]. These antibodies were originally isolated for evaluation as potential therapeutic interventions in SARS-CoV-2 infected patients. In this current study we have used RBD-specific antibodies in this hMAb panel to examine the cellular properties of the S protein RBD in virus-infected African green monkey kidney epithelial cells (Vero E6 cells). Our experimental approach is aimed to complement the ongoing high-resolution structural studies undertaken by other groups on the S protein and the interaction of the RBD with neutralising antibodies [22, 23]. Our current analysis was performed using SARS-CoV-2 clinical isolates that were isolated during the early phase of the pandemic in Singapore. These immunological reagents also allowed us to perform a comparison of the biological properties of these SARS-CoV-2 clinical isolates in infected cells.

## Materials and Methods

### Isolation of SARS-CoV-2

Three isolates of SARS-CoV-2 viruses (designated as 1302, 0563 and 0334) were isolated from nasopharyngeal samples during the early COVID 19 pandemic in Singapore. These strains were first detected using PCR primers and thermocycling conditions described in [24]. DSO’s Institutional Ethics Review Board (IRB no 0008/2020) approval was sought before using the de-identified nasopharyngeal samples for virus isolation. The SARS-CoV-2 viruses were isolated in the African green monkey kidney epithelial (Vero E6) cells (CCL-81, American Type Culture Collection, Virginia, USA) and maintained in Minimum Essential Medium (MEM) supplemented with 10% fetal bovine serum (FBS) (Invitrogen, Carlsbad, California, USA) and 1% penicillin and streptomycin (Invitrogen). Virus infection was carried out in the same medium with 2 % FBS at 37^0^C in the presence of 5% CO_2_. The infectivity was titrated using standard virological protocols described in [19]. In Singapore, SARS-CoV-2 is classified as Risk Group 3 and this is governed by the Biological Agent and Toxins Act (https://www.moh.gov.sg/biosafety/about-bata assessed on 26th November 2021). Hence, all the experiments with live SARS-CoV-2, including primary isolation were conducted in the biosafety level 3 laboratory located at the DSO National Laboratories, with protocols (Protocol numbers BSL3-2020000001 and BSL3-2020000008) approved by DSO’s Institutional Biosafety Committee and the Ministry of Health.

### Amplification and sequencing of S gene

Total nucleic acid extraction was performed on the different virus isolates and passages 1 to 3, using the QIAamp DNA mini-kit (Qiagen, Hilden, Germany) according to the manufacturer’s instructions. For amplification of the S gene, reverse transcription was first performed on the extracted nucleic acid with random hexamer primer using the first strand cDNA synthesis kit (Thermo Fisher Scientific, Waltham, Massacchusetts, USA) according to the manufacturer’s instructions. The 3.8 kbp S gene was PCR amplified with primer SARS-CoV-2-S-F (5’- ATG TTT GTT TTT CTT GTT TTA TTG C-3’) and primer SARS-CoV-2-S-R (5’- TTA TGT GTA ATG TAA TTT GAC TCC T-3’) using the PCR reaction mix consisting of 1× PCR buffer, 0·5 μM primers, 0.1 mM dNTPs, 1.0 U Q5® High Fidelity DNA polymerase (New England Biolabs, Ipswich, Massacchusetts, USA) and 5 μl of viral RNA in a final volume of 50 μl. The following thermocycling conditions were used: an initial denaturation at 98 °C for 30 s, followed by 35 cycles at 98 °C for 10 s, 59 °C for 30 s, and 72 °C for 4 min with a final extension step at 72 °C for 2 min. The PCR products were electrophoresed on 1% agarose in 0.5x Tris-borate-EDTA (TBE) buffer and visualized by Midori Green (Nippon Genetics Europe GmbH, Düren, Germany) staining. The 3.8 kbp amplicons were sent for Sanger sequencing. The nucleotide or amino acid sequences were assembled and aligned with the T-Coffee program which used both the Clustal W and Lalign methods for multiple sequence alignment [25]. The full-length S gene sequences of different virus isolates and passages were next translated to amino acid sequence using the Expert Protein Analysis System (Expasy) from Swiss Bioinformatic Resource Portal (https://www.expasy.org/; [26]). Multiple-aligned amino acid sequences of S proteins were generated using the Boxshade program of Expasy.

### Discovery and isolation of human monoclonal antibodies

SC23 and SC29 were generated by single B cell antibody interrogation from convalescent patient sample as described in [19]. The human monoclonal antibodies, PD4, PD5 and PD7 were isolated from an immune phage display library constructed from the convalescent patients’ B cells according to the methods by [27]. PD4, PD5 and PD7 were obtained by biopanning against recombinant expressed SARS-CoV-2 S protein using the method described in [19].

### Antibodies and specific reagents

The rabbit polyclonal antibody to S protein (polyS) (Sino Biological, Singapore) and anti-N (Thermo Fisher Scientific) were purchased. The anti-rabbit, anti-mouse and anti-human IgG conjugated to Alexa 488 and Alexa 555, and wheat germ agglutinin conjugated to Alexa 488 were purchased from Invitrogen (Thermo Fisher Scientific). The giantin rabbit polyclonal antibody was obtained from Dr Lu Lei (NTU). Publicly available sequences of the Regeneron antibodies casirivimab and imdevimab (REGN-10933 and REGN-10987) were transiently expressed in HEK293 suspension culture and purified by Protein A affinity chromatography on FPLC as described previously [19]. The furin inhibitor decanoyl-RVKR-cmk and mannosidase-1 inhibitor deoxymannojirmycin (DMJ) were purchased from Calbiochem (San Diego, California, USA).

### Enzyme Linked Immunosorbent Assay (ELISA)

Recombinant Spike extracellular domain (WT) and RBD proteins were expressed and purified through Twin-Strep tag for use in ELISA. 2 µg/ml of purified protein was diluted in Binding buffer (100 mM Tris-HCl, 1 mM EDTA, 150 mM NaCl, pH 8.0) and coated onto StrepTactin XT 96-well ELISA plates, 100 µl/well, for 2 hr at room temperature (RT) before washing twice with PBS. Antibodies were diluted in blocking solution (2% BSA/PBS) to the indicated concentrations and added to the coated plate at 100 µl/well and incubated for 1hr at RT, before washing thrice with PBS/0.05%Tween. Antibody binding was detected using goat anti-Human IgG Fc Secondary Antibody conjugated to horse radish peroxidase (Thermo Fisher Scientific), diluted 1:5000 in blocking solution and incubated for 1 hr at RT. Plates were washed thrice with PBS/0.05% Tween and once with PBS. After washing, plates were developed with tetramethylbenzidine (TMB) substrate (Thermo Fisher Scientific). The reaction was stopped with 2 M sulphuric acid and absorbance was measured at 450 nm.

### Neutralization assay

The microneutralization assay was performed as described in [19]. Briefly, antibodies at indicated concentrations was incubated with 100 TCID_50_ of hCoV-19/Singapore/3/2020 virus and 2 x 10^4^ Vero E6 cells in 100 μl of culture media in 96-well flat bottom plates and incubated for 72 hrs. The neutralization was measured using Viral Toxglo reagent (Promega, Madison, Wisconsin, USA) to determine the percentage of cell survival relative to uninfected and virus only controls.

### Virus infection

Vero E6 cells were seeded onto 12 mm circular glass coverslips and infected with SARS-CoV-2 at the required multiplicity of infection at 37°C. When appropriate, 40µM decanoyl-RVKR-cmk was added from 4 hr post-infection. At the required time the cells were washed with PBS and fixed with 4% paraformaldehyde (Sigma-Aldrich, Burlington, Massacchusetts, USA) in PBS for 30min prior to further processing.

### Lactate Dehydrogenase (LDH) cytotoxicity assay

Cell cytotoxicity was assessed by measuring LDH release from cells using the LDH cytotoxicity assay (Promega). This was performed on non-treated and decanoyl-RVKR-cmk-treated Vero cells following the manufacturer’s instructions.

### Recombinant S protein expression

The S gene of the SARS-CoV-2 (Accession No. MN908947) was synthesized by Twist BioSciences (San Francisco, USA). The codon-optimized full-length S gene (nucleotide residues 1 to 1273) was assembled as previously described [19] and cloned into the recombinant pCAGGS vector to create pCAGGS/S. For S1 subunit and RBD expression, only residues 1-685 (S1) and 331-524 (RBD) were cloned together with a c-flag and a c-myc tag respectively. The fragments were cloned into pCAGGS to generate pCAGGS/S1 and pCAGGS/RBD. Bulk preparation of all plasmids was performed using the plasmid midiprep kit (Qiagen). Cells (1×10^5^) were transfected into HEK293 with 1µg plasmid DNA using Lipofectamine 2000 (Invitrogen) following the manufacturer’s instructions. The transfected cells were maintained at 37°C until the time of sample processing.

### Immunofluorescence microscopy

The cells on 12 mm circular glass coverslips were fixed with 4% paraformaldehyde (Sigma-Aldrich) and washed with PBS. The cells were either non-permeabilized, or permeabilized using 0.1% Triton X100 in PBS at 4°C for 15 mins prior to antibody staining. The cells were stained with the appropriate primary and secondary antibody combinations and mounted on microscope slides using Citifluor. The stained cells were imaged using a Nikon Eclipse 80i Microscope (Nikon Corporation, Tokyo, Japan) and an Etiga 2000R camera (Q Imaging, Teledyne Photometrics, Tucso Arizona, USA) attached. The images of immunofluorescence-stained cells were recorded using Q Capture Pro ver 5.0.1.26 (Q Imaging, Teledyne Photometrics). Imaging for confocal microscopy was performed with a Zeiss 710 confocal microscope (Zeiss, Oberkochen, Germany) with Airyscan using appropriate machine settings. The images recorded were examined and processed using Zen ver 2.3 software (Zeiss).

### Scanning electron microscopy (SEM)

The cells on 13 mm glass coverslips were fixed using 4% paraformaldehyde (Sigma-Aldrich) in PBS and washed with PBS. The cells were then fixed sequentially using 3% glutaraldehyde (Sigma-Aldrich) and 1% Osmium tetroxide (Sigma-Aldrich), dehydrated using an ethanol gradient and critical point drying performed as described previously [28]. The processed cells were gold coated and mounted on aluminium stubs and imaged with a FEI QUANTA 650 FEG scanning electron microscope (FEI company, Oregon, USA) using appropriate machine settings.

## Results and discussion

### 1. The Singapore SARS-CoV-2 isolates and the human monoclonal antibody panel used in this study

In this study we used three SARS-CoV-2 viruses that were isolated during the early phase of COVID19 pandemic in Singapore, which are referred to as SARS-CoV-2/1302, SARS-CoV-2/0563 and SARS-CoV-2/0334. Since the characterization of the S protein hMAbs was the major focus of this study, the complete S protein sequence for each isolate used in our analysis was determined. Each virus isolate was passaged three times in Vero E6 cells and the genetic material extracted from each passage was PCR-amplified with specific primers for the S gene and sequenced using Sanger’s sequencing. The sequences from overlapping amplicons were assembled and analyzed. The data revealed that the nucleotide and amino acid sequences of the S protein from the three Singapore isolates had a high level of sequence homology to the S sequence of SARS-CoV-2/WIV04 isolate (GenBank Accession MN996528) that was reported when the COVID 19 pneumonia first originated in Wuhan in China [7]. In the first passage the S protein sequence of each virus isolate was identical to the S protein sequence SARS-CoV-2/WIV04 isolate (SFig. 1). However, the subsequent passages of the virus during virus stock amplification showed some specific changes in the primary amino acid sequences of the S protein. The S protein sequence of the SARS-CoV-2/0334 and SARS-CoV-2/0563 comprised of 1269 amino acids and were 100% identical. Each had a five amino acid deletion in the S protein sequence at _675_QTQTN_679_ that was just up-stream of the polybasic furin cleavage site (SFig 1). Although the _675_QTQTN_679_ deletion was not observed in the S protein sequence of SARS-CoV-2/1302, the S protein sequence of SARS-CoV-2/1302 had a single amino acid change R682W in the furin cleavage site. This created _682_WRARS_686_ rather than the complete furin consensus sequence _682_RRARS_686_ that is found in the S protein sequence of SARS-CoV-2/WIV04, SARS-CoV-2/0563 and SARS-CoV-2/0334. Virus variants, also known as quasi-species, containing mutations in the region of the S1/S2 protein proteolytic cleavage site have been reported to be present in SARS-CoV-2-infected patients [29], and these can be selected during passaging of the virus in tissue culture. The basis for this virus selection and the selective advantage that these sequence changes impart to the virus isolate in tissue culture is currently unclear. Apart from the sequence differences indicated above, the remaining S protein sequence of each of the Singapore isolates was 100% identical to the S protein sequence of SARS-CoV-2/WIV04. In particular, the sequence of the S protein RBD was 100% identical in SARS-CoV-2/WIV04 and all three of the Singapore SARS-CoV-2 isolates. Since the S protein sequences of the SARS-CoV-2/0563 and SARS-CoV-2/0334 isolates were identical, all subsequent work on characterizing immune-reactivity of the hMAb panel was performed mainly using the SARS-CoV-2/1302 and SARS-CoV-2/0334 isolates.

In this this study we selected PD4, PD5, PD7, SC23 and SC29 from the original antibody panel for further characterisation using a cellular virology approach. We failed to detect binding of these antibodies to the S protein using Western blotting (SFig. 2A), but recognition of S protein by each antibody in the panel was confirmed by using the purified SARS-CoV-2 S protein ectodomain in ELISA (Fig. 1A). In a similar analysis using a sequence that corresponded to the S protein RBD, only PD5, PD7 and SC23 showed RBD binding (RBD-binders) (Fig. 1B(i)). It is presumed that the epitopes recognized by the RBD-binders are formed by different parts of the RBD once it had folded into the correct conformation. This folding pattern is expected to be lost during the sample processing process in the Western blot analysis. The inability of these antibodies to bind to the S protein in Western blot analysis provided evidence that the RBD-binders bind to conformational-specific epitopes on the correctly folded S protein. Although comparable binding affinities to the complete S protein was noted for all hMAbs, the SC23 showed reduced binding-affinity for the RBD compared with PD5 or PD7 (Figs. 1B(ii) and 1C). This indicate that SC23 may recognize a distinct sequence in the RBD and that one or more sequences located outside of the RBD may facilitate its binding to the RBD. No RBD-binding was detected by PD4 and SC29, suggesting that the SC29 and PD4 epitope recognition sequences were outside the RBD region. Although we have not mapped the binding domains of PD4 and SC29, we examined their binding activities using an ELISA-based antibody-binding competition assay (SFig. 2B). This assay indicated that the binding of PD4 and SC29 antibodies to the S protein was mutually exclusive, suggesting that PD4 and SC29 bind to similar locations on the S protein. Prior binding of either the PD5, PD7 or SC23 antibodies to the S protein did not interfere with PD4 or SC29 binding, which was consistent with PD4 and SC29 being non-RBD binders (SFig. 2B). Binding of the PD5, PD7 and SC23 antibodies to the S protein in the same antibody-binding competition assay were also mutually exclusive indicating binding to similar locations on the S protein and consistent with their RBD-binding properties. There was a general correlation between the RBD recognition and the complementarity-determining regions (CDRs) sequences of the individual antibodies in the hMAb panel. There was a high degree of similarity between the sequence of the CDRs of PD5 and PD7 and to a lesser extent with SC23 (SFig. 2C), which correlated with the different RBD binding activity of SC23. The PD4 and SC29 showed high levels of sequence similarity in the CDR1 and CDR2 in the heavy chain and was distinct from the CDRs in the RBD-binders.

**Fig. 1.**
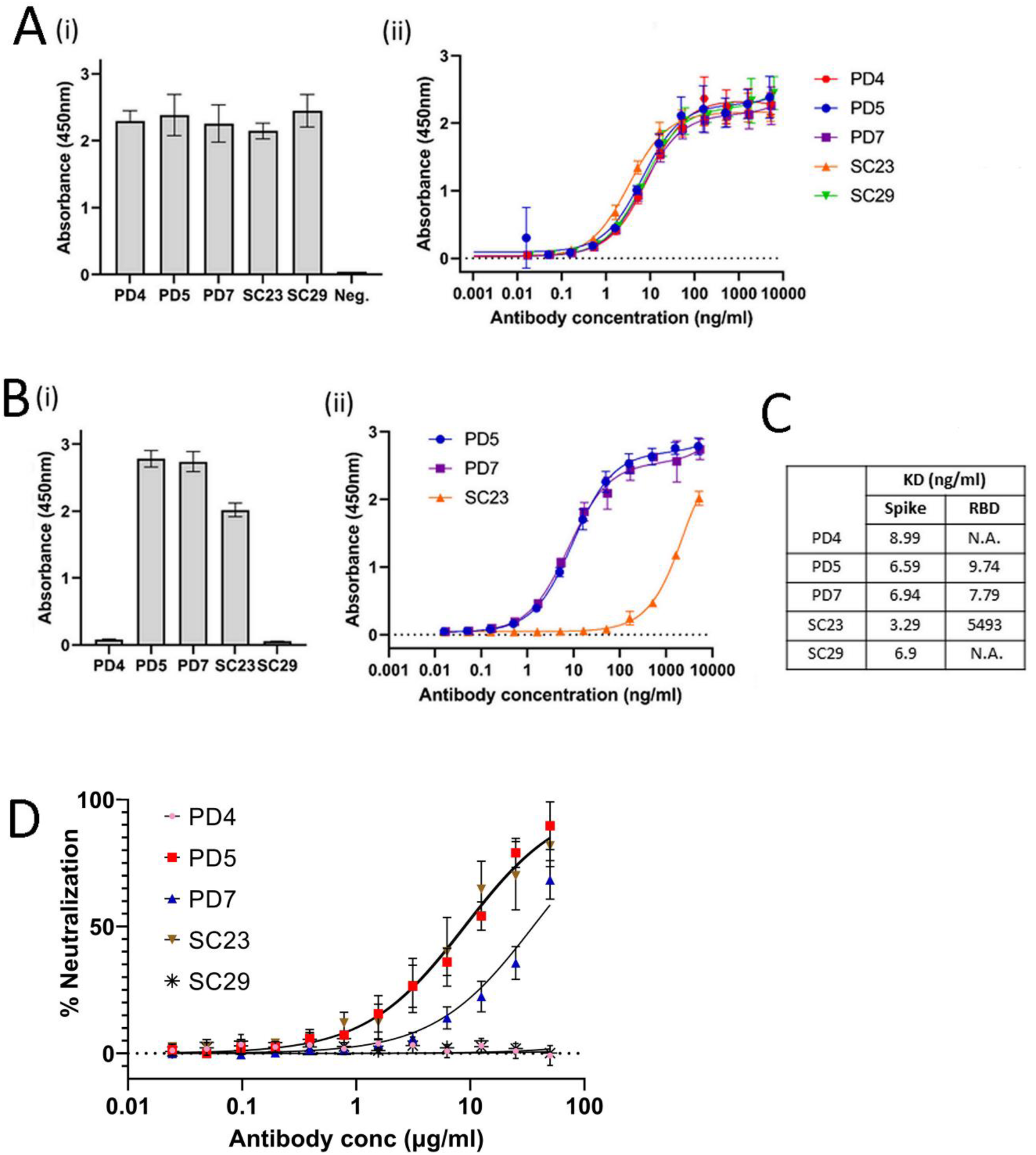
Specificity of anti-SARS-Cov-2 human monoclonal antibodies used in this study. The binding affinities to the spike (S) protein and neutralization activity with SARS-CoV-2 virus was determined for each human monoclonal antibody (hMAb) in the panel, PD4, PD5, PD7, SC23, SC29. **(A)** (i) Binding affinity of the hMAb panel against the extracellular domain of the S protein by enzyme-linked immunosorbent assay (ELISA); (ii) Binding affinity of the hMAb panel to the S protein as a function of antibody concentration from 0.001 to 10,000 ng/ml. **(B)** (i) Binding of hMAb panel to the receptor binding domain (RBD) of the S protein as measured by ELISA assay; (ii) Binding of the PD5, PD7 and SC23 to the RBD of S protein as a function of antibody concentration from 0.001 to 10,000 ng/ml. **(C)** The binding affinities to the extracellular domain and the RBD of S protein was determined for each member of the hMAb panel and measured in the dissociation constant (KD in ng/ml). (NA; not applicable). **(D)** Neutralization activity of the hMAb panel with 100 TCID_50_ of infectious SARS-CoV-2 and measured as a function of antibody concentration (Antibody conc) from 0.01 to 100 µg/ml. Neutralization efficacy is represented as a percentage relative to uninfected and virus only controls.

The differential recognition of the RBD was further supported by examining the ability of the hMAb panel to block SARS-CoV-2 binding to Vero E6 cells using a virus neutralization assay. Only PD5, PD7 and SC23 exhibited virus neutralizing activity, while PD4 and SC29 failed to inhibit infection (Fig. 1D). These data indicated that inhibition of SARS-CoV-2 infection correlated with RBD recognition, and it is presumed that binding of the antibodies to the RBD would create a steric hindrance that would interfere with the virus attachment to the cell receptor.

### 2. Distribution of the S protein RBD in SARS-CoV-2-infected Vero E6 cells

The recognition of S protein by each antibody in hMAb panel was further confirmed by examining their immuno-reactivity in SARS-CoV-2-infected Vero E6 cells (Fig. 2A) Vero E6 cells were either mock-infected or infected with SARS-CoV-2/0334 and at 18 hpi the cells were stained using either PD5, PD7, PD4, SC23, and SC29 (Fig. 2A). Imaging using IF microscopy showed that fluorescence staining using either antibody was only detected in the virus-infected cells, indicating S protein recognition by the hMAb panel. Virus infection was confirmed by using the commercially available S protein polyclonal antibody (polyS) that recognizes the S2 domain and the N protein antibody (anti-NP).

**Fig. 2.**
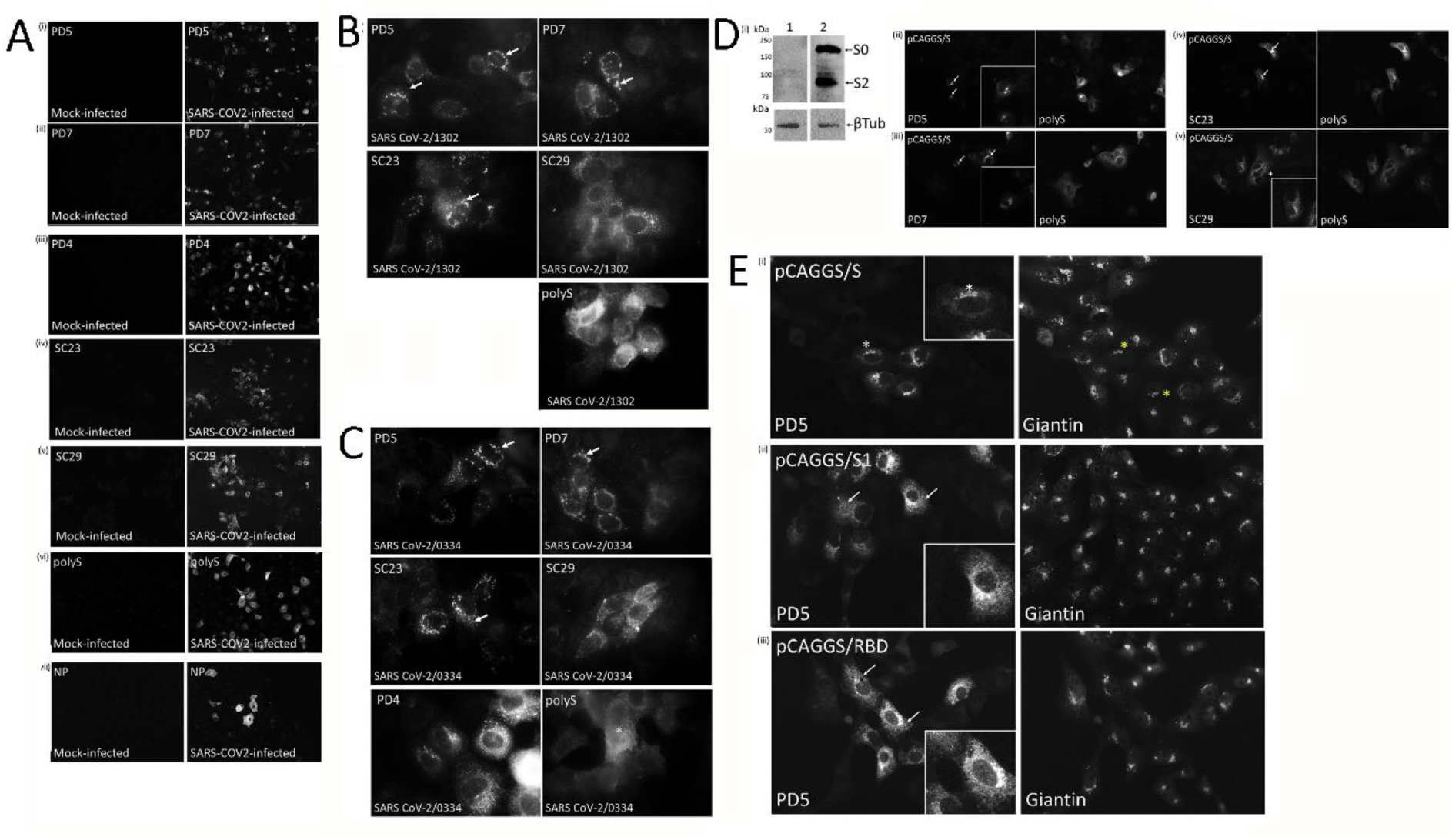
Immune reactivity of the anti-SARS-CoV-2 human monoclonal antibodies in virus-infected Vero E6 cells. **(A)** At 18 hrs post-infection (hpi), mock-infected and SARS-CoV-2/0334-infected cells were stained using (i) PD5, (ii) PD7, (iii) PD4, (iv) SC23, (v) SC29, (vi) polyS and (vii) anti-NP, and imaged using immunofluorescence (IF) microscopy (objective x40 magnification). Vero E6 cells were infected with **(B)** SARS-CoV-2/1302 and **(C)** SARS-CoV-2/0334, and at 18 hpi the cells were co-stained using PD5, PD7, PD4, SC23, SC29 and polyS as indicated. The stained cells were imaged using IF microscopy ((objective x100 magnification; oil immersion). The prominent punctate cytoplasmic staining pattern (white arrow) is highlighted. **(D)** (i) Cells were (1) mock-transfected and transfected with (2) pCAGGS/S and the lysates immunoblotted with polyS. Protein species corresponding in size to the uncleaved (S0) and S2 domain of the S protein are indicated. Tubulin is the loading control. Cells were transfected with pCAGGS/S and co-stained with polyS and (ii) PD5, (iii) PD7, (iv) SC23 and (v) SC29 as indicated. The stained cells were imaged by IF microscopy (objective x40 magnification). The diffuse cytoplasmic S protein staining (*) and punctate cytoplasmic staining (white arrows) are indicated. **(E)** Cells were transfected with (i) pCAGGS/S, (ii) pCAGGS/S1 and (iii) pCAGGS/RBD as indicated and co-stained with PD5 and anti-giantin, The co-stained cells were imaged by IF microscopy (objective x40 magnification). Inset is an enlarged image of a representative cell showing the specific PD5 staining in each case.

Cells infected with either the SARS-CoV-2/1302 (Fig. 2B) or SARS-CoV-2/0334 (Fig. 2C), were stained with PD5, PD7, PD4, SC23, SC29 and polyS and examined in greater detail using IF microscopy. In SARS-CoV-2 infected cells stained with PD5, PD7 and SC23, a similar prominent punctate cytoplasmic staining pattern was apparent, while cells stained with either PD4 or SC29 exhibited a more distinct cytoplasmic staining pattern. These staining patterns were defined as being representative if they were observed in greater than 94% of the cells in the field of view using IF microscopy (e.g. at 20x magnification), and in several replicate experiments (i.e. greater than 4 individual experiments). The diffuse PD4 or SC29 staining pattern was similar in appearance to the staining pattern exhibited by polyS-stained virus-infected cells. The imaging data therefore indicated that the hMAb panel could be divided into two groups based on their staining pattern in virus-infected cells; a prominent localized punctate staining pattern (PD5, PD7 and SC23) or a more broadly diffuse cytoplasmic staining pattern, (PD4 and SC29), and these antibody-staining patterns correlated with RBD-recognition. These data suggested that the RBD-binders (e.g. PD5) may recognize a distinct population of the S protein which allows recognition of the RBD. Since the polyS is a polyclonal S protein antibody, it would be expected to detect different forms of the S protein in virus-infected cells, albeit by that recognizing the S2 domain.

In non-infected Vero E6 cells expressing the recombinant S protein cell lysates were prepared and examined by immunoblotting with polyS (Fig. 2D(i)). S protein species corresponding in size to the uncleaved S protein (S0) and to the S2 domain were detected, which confirmed that the recombinant S protein was correctly processed in the transfected cells. The recognition of the S protein in the immunoblotting assay by polyS also suggested that recognition of the S protein by this antibody involved one or more of the liner epitopes rather than conformational specific epitopes in the hMab panel. Cells expressing the recombinant S protein were co-stained with polyS and either PD5, PD7, SC23 and SC29 and examined using IF microscopy (Fig. 2D (ii)-(v)) and staining of the transfected cells with either antibody confirmed the S protein recognition. We also compared the PD5 staining in cells expressing the full-length recombinant S protein with those expressing either only the S1 domain or the RBD (Fig. 2E). A more widespread PD5 staining pattern was observed in cells expressing the S1 domain and the RBD compared with the cell expressing the full-length S protein sequence. This confirmed that the RBD can form into its distinct structure independently of the S2 domain and indicated that the PD5 can recognize the RBD at other locations in cells expressing the S protein. This further supports the suggestion that in SARS-CoV-2-infected cells the PD5 may recognize a distinct population of the S protein in which the RBD is accessible to antibody binding.

The IF microscopy analysis described above allowed several antibody stained cells to be imaged in the same field of view, allowing representative antibody staining patterns to be determined. In contrast, confocal microscopy allows the detailed imaging of the antibody staining in individual representative cells. Since the respective antibody staining patterns were similar for both SARS-CoV-2 isolates, we used confocal microscopy to examine the staining pattern for each antibody in SARS-CoV-2/0334-infected Vero E6 cells. At 18 hpi the cells were co-stained with polyS and either PD5, PD7, PD4, SC23 and SC29, and a series of images was recorded in the Z-plane from individual representative co-stained cells (SFig. 3). Single images were extracted at an optical plane from the Z-series that allowed the cytoplasmic staining patterns of the different antibodies to be compared (Fig 3A). The polyS antibody exhibited a diffuse cytoplasmic staining pattern, but a small degree of co-localization within the distinct prominent punctate staining pattern exhibited by PD5, PD7 and SC23 was noted. This punctate staining pattern was also exhibited in PD4- and SC29-stained virus-infected cells, however these antibodies also exhibited an additional prominent diffuse SC29- and PD4- staining patterns. These different antibody staining patterns suggested that the PD5, PD7 and SC23 only recognize a specific subpopulation of the total S protein in which the protein conformation renders the RBD accessible to antibody binding. The recognition of the S protein by PD4 and SC29 was not dependent on RBD recognition, and the additional diffuse antibody staining pattern suggested that they recognize other forms of the S protein. A similar imaging analysis of transfected cells expressing the recombinant S protein and co-stained with polyS and PD5 and SC29 was also performed. This showed the distinct antibody-specific staining patterns that were similar to that observed in SARS-CoV-2-infected cells (Fig. 3B) in which the PD5 staining exhibited a general localized staining pattern compared with the more widespread staining pattern of SC29. This is consistent with similar processing of the recombinant S protein and the S protein expressed in virus-infected cells and indicated that the different S protein antibody staining patterns that we observed was not dependent on virus infection.

**Fig. 3.**
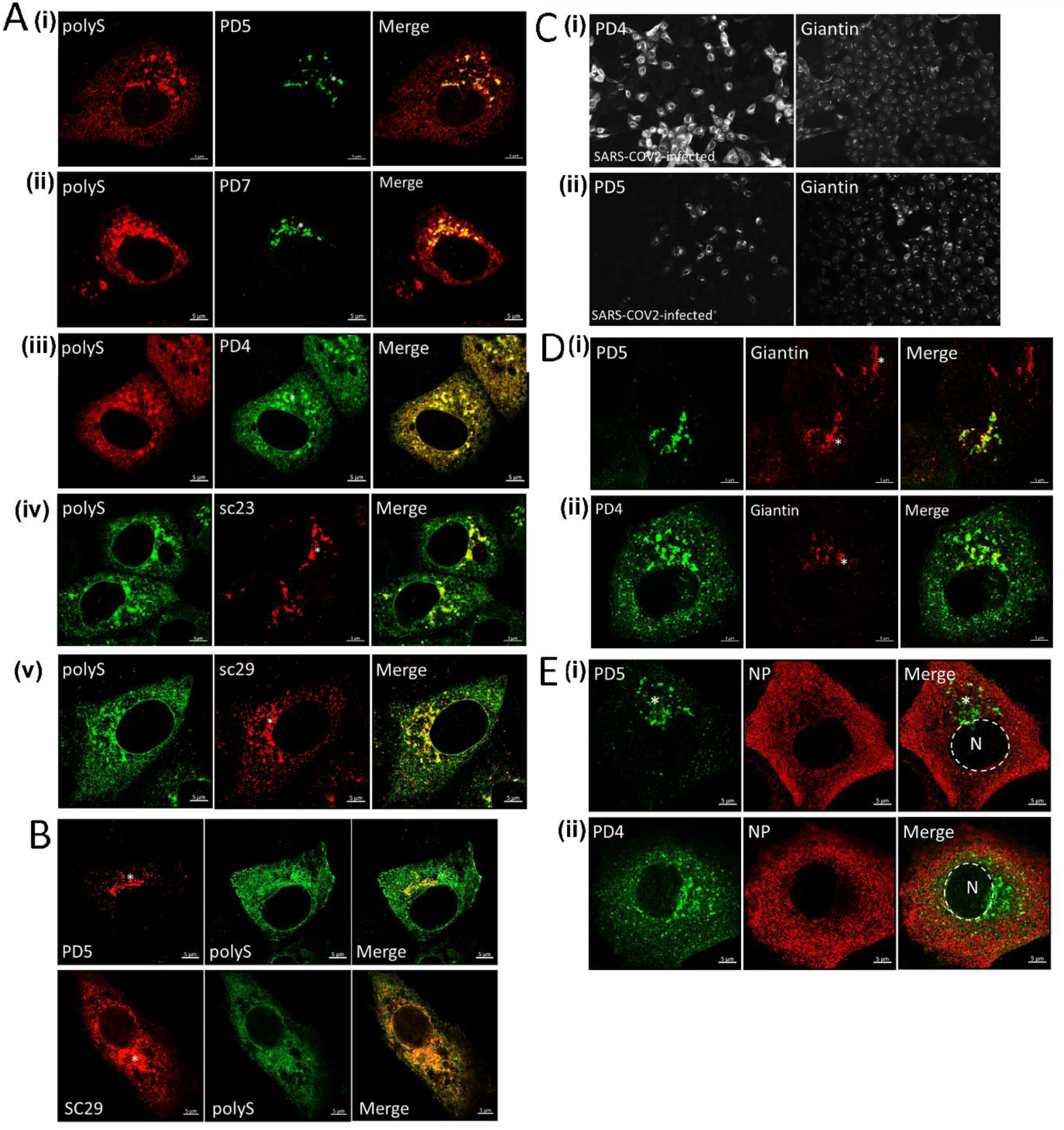
Distribution of the S protein RBD and Golgi complex in virus-infected Vero E6 cells. **(A)** At 18 hrs post-infection SARS-CoV-2/0334-infected cells were co-stained using anti-polyS and either (i) PD5, (ii) PD7, (iii) PD4, (iv) SC23 and (v) SC29 and imaged using confocal microscopy. The individual channel and merged images are shown. The prominent punctate cytoplasmic staining pattern (*) is highlighted. **(B)** Vero E6 cells expressing recombinant S protein were co-stained with anti-polyS and either PD5 or SC29 and imaged using confocal microscopy. The individual channel and merged images are shown. **(C)** SARS-CoV-2/0334-infected cells were co-stained using (i) PD4 and anti-giantin and (ii) PD5 and anti-giantin and imaged using immunofluorescence microscopy (objective x20 magnification;). **(D)** SARS-CoV-2/0334-infected cells were co-stained using (i) PD5 and anti-giantin and (ii) PD4 and anti-giantin and imaged using confocal microscopy The giantin staining pattern indicating the Golgi complex (*) is highlighted. **(E**) SARS-CoV-2/0334-infected cells were co-stained using (i) PD5 and anti-NP and (ii) PD4 and anti-NP and imaged using confocal microscopy. The punctate PD5 staining pattern (*) and area of the nucleus (N and delineated by the white broken line) are indicated. In all confocal microscope images the individual channel images and merged images are shown..

The C-terminus of the S protein contains an ER retrieval signal which facilitates the accumulation of the S protein close to the site of coronavirus particle assembly [30]. The process of coronavirus assembly involves the ER-Golgi intermediate compartment (ERGIC) and Golgi complex [6]. The S protein undergoes a series of important post-translational modifications that are mediated by cellular activities associated with the Golgi complex (e.g. glycosylation, furin cleavage) and in virus-infected cells we examined the staining patterns of the RBD-binders in the context of the Golgi complex. The PD5, PD7 and SC23 exhibited similar staining patterns within infected cells, and we concluded that imaging of PD5 staining could be used as a representative of this antibody group (RBD-binders). Mock and SARS-CoV-2/0334-infected Vero E6 cells were co-stained with PD5 or PD4 and anti-giantin (a Golgi complex marker) and imaged using IF microscopy. Although some co-staining between PD4 and anti-giantin was noted, the overall diffuse PD4 staining pattern contrasted with the localized anti-giantin staining (Fig. 3C(i)), Similar PD5 and anti-giantin staining pattern in virus-infected cells was noted (Fig. 3C(ii)), indicating that PD5 staining was largely localized to elements of the Golgi complex. A more detailed analysis of the antibody staining patterns was performed using confocal microscopy on PD5 and PD4 cells contained with anti-giantin (Fig. 3D). This revealed a high level of co-localization between the punctate PD5 staining pattern and the Golgi complex in infected cells (Fig. 3D(ii) and (iii)) and SFig 3F) and confirmed that the punctate PD5 staining pattern was mainly restricted to the Golgi complex. The PD4 staining exhibited the diffuse antibody staining described above which only partially localized with the anti-giantin staining at the Golgi complex (Fig. 3D(ii)). The infected cells were also co-stained with either PD5 or PD4 and anti-NP and imaged using confocal microscopy (Fig 3E). The anti-NP labelling gives rise to a prominent cytoplasmic staining pattern that allowed delineation of the infected cells and the absence of nuclei staining with this antibody enables the position of the nucleus to be visualized. This staining combination therefore allows the respective PD5 and PD4 staining pattern to be visualized in the context of the whole cell.

We also examined staining pattern of REGEN-10933 and REGEN-10987 whose binding sites in the S protein RBD have been accurately defined using structural biology [31, 32]. SARS-CoV-2/0334 infected cells were co-stained with either REGEN-10987 or REGEN-10933 and anti-NP (SFig. 4). The anti-NP gave rise to a broadly cytoplasmic staining pattern that delineated the body of the cell, which contrasted with the localized and punctate staining pattern exhibited by REGEN-10933 and REGEN-10987. The REGEN-10933 and REGEN-10987 antibodies also showed staining patterns that was similar to the staining patterns exhibited by the RBD-binders in virus-infected cells.

These data suggested that RBD recognition by PD5 occurs as the S protein is trafficked through the Golgi complex. The reason for the apparent accumulation of the S protein at the Golgi complex during its transport through the secretory pathway is currently unclear, but it may be related to the extensive glycosylation of the S protein [33]. Although virus assembly is proposed to occur close to the Golgi complex, similar PD5-staining pattern was also observed in transfected cells expressing the recombinant S protein, suggesting that this Golgi-localisation mays not directly caused by virus induced changes in the secretory pathway.

The imaging data described above demonstrated the specificity of the hMAb panel with regards to the S protein recognition in virus-infected Vero E6 cells and demonstrated specific antibody staining patterns that correlated with RBD-binding. Furthermore, these data also indicated that the deleted sequence _675_QTQTN_679_ in SARS-CoV-2/0334 and the R682W change in SARS-CoV-2/1302 (SFig.1) did not prevent recognition by either of the antibodies examined. We have failed to detect PD5 co-staining with antibodies against early compartments of the secretory pathway e.g. the ER compartment (Sugrue unpublished observations), suggesting that PD5 staining is only detected in the later compartments of the secretory pathway. The more widespread cytoplasmic staining of the non-RBD binders and polyS is presumably due to recognition of the S protein at other cellular locations (in addition to the Golgi-complex). These data therefore suggest that while the non-RBD binders and polyS recognise the total S protein expressed in virus-infected cells, the RBD-binders recognise only a subset of the total S protein where the RBD is antibody accessible. Although the Golgi compartment may be modified in SARS-CoV-2-infected cells to facilitate virus replication, the staining pattern of the RBD-binders (exemplified by PD5) was mainly associated with the Golgi-complex. Since the PD5 recognises conformational specific epitopes within the RBD of the S protein, these data suggest that a conformational change in the S protein may occur that leads to the correctly folded RBD at the Golgi complex that enabled PD5 recognition.

### 3. The human monoclonal antibodies recognize the RBD displayed on virus particles on the surface of SARS-CoV-2/1302 and SARS-CoV-2/0334 infected Vero E6 cells

We used imaging to examine the surface expression of the S protein on infected Vero E6 cells stained with each of the antibodies in the hMAb panel. Vero E6 cells were mock-infected (Fig. 4A) and infected with either SARS-CoV-2/0334 (Fig. 4B(i)) or SARS-CoV-2/1302 (Fig. 4B(ii)), and at 18 hpi the non-permeabilized cells were co-stained using PD5 and polyS and imaged using IF microscopy. A similar diffuse surface-staining pattern was noted on cells infected with either virus isolate. A similar surface staining pattern was also observed on non-permeabilized cells infected with SARS-CoV-2/0334 and co-stained with polyS and either PD7, SC23 and SC29 (Fig. 4C). The surface staining with the PD5, PD7 and SC23 antibodies indicated that the RBD was displayed on the surface of infected cells. Although SC29 did not recognize the RBD and did not exhibit virus neutralization activity, surface staining on SC29-stained non-permeabilized cells indicated that the SC29-epitopes were also surface-displayed.

**Fig. 4.**
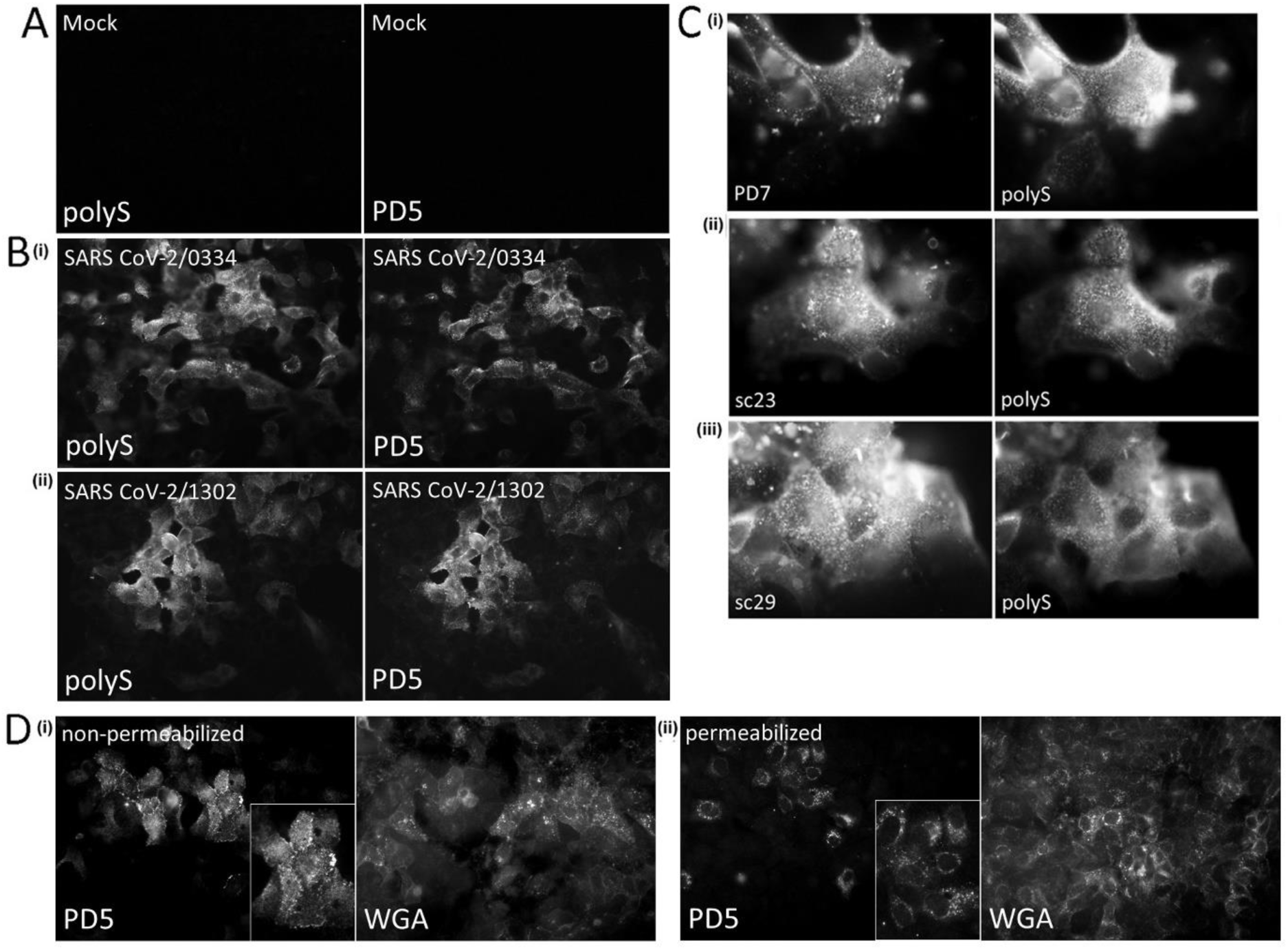
Surface staining of SARS-CoV-2 infected Vero E6 cells. Vero E6 cells were **(A)** mock-infected and **(B)** infected with the (i) SARS-CoV-2/0334 or (ii) SARS-CoV-2/1302. At 18 hrs post-infection (hpi) the non-permeabilized cells were co-stained using polyS and PD5. The stained cells were imaged using immunofluorescence (IF) microscopy (objective x40 magnification). **(C)** Non-permeabilized SARS-CoV-2/0334-infected cells were co-stained with polyS and either (i) PD7, (ii) SC23 or (iii) SC29. The stained cells were imaged using IF microscopy (objective x100 magnification; oil immersion). **(D)** At 18 hpi SARS-CoV-2/0334-infected cells were (i) non-permeabilized and (ii) permeabilized, and the cells co-stained using PD5 and wheat germ agglutinin conjugated to Alexa 488 (WGA). The stained cells were imaged using IF microscopy (objective x40 magnification). In both (i) and (ii) the inset is an enlarged imaged showing the PD5 staining pattern in each condition.

The surface staining patterns were confirmed by comparing the PD5-staining patterns of non-permeabilized cells and permeabilized SARS-CoV-2/0334-infected cell. Wheat germ agglutinin conjugated to Alexa 488 (WGA-488) is an established cellular probe that stains cellular membranes and cells and were co-stained with PD5 and WGA-488 and imaged by IF microscopy (Fig. 4D). As expected, the PD5-staining pattern observed on non-permeabilized cells (Fig. 4D (i)) was clearly distinct from the PD5 punctate cytoplasmic staining pattern exhibited on permeabilized cells (Fig. 4D (ii)). The non-permeabilized SARS-CoV-2/0334-infected cells were co-stained using PD5 and polyS and examined in greater detail using confocal microscopy. A series of images was recorded in the Z-plane from individual representative co-stained cells which allowed surface staining at the cell top (Fig. 5A(i)) and at the cell periphery to be imaged (Fig. 5A(ii)). Both antibodies exhibited a similar surface staining pattern on the co-stained cells, which appeared as small spots and filament, and was distinct from the cytoplasmic PD5 and polyS staining patterns in permeabilized cells described above. The level of co-localization of these antibodies was examined using the Pearson’s and Mander’s correlation coefficients (Fig. 5B), which indicated a high level of co-localization between the two antibodies. A high level of co-staining between both antibodies would be expected since both antibodies would be expected to recognize the same population of the S protein on the surface of infected cells. The surface-staining pattern exhibited by both antibodies was more apparent in a 3-D reconstruction of the permeabilized co-stained infected cell (Fig. 5C (i) and (ii))), where the filamentous co-staining pattern was clearly distinguished from the spotted antibody staining pattern.

**Fig. 5.**
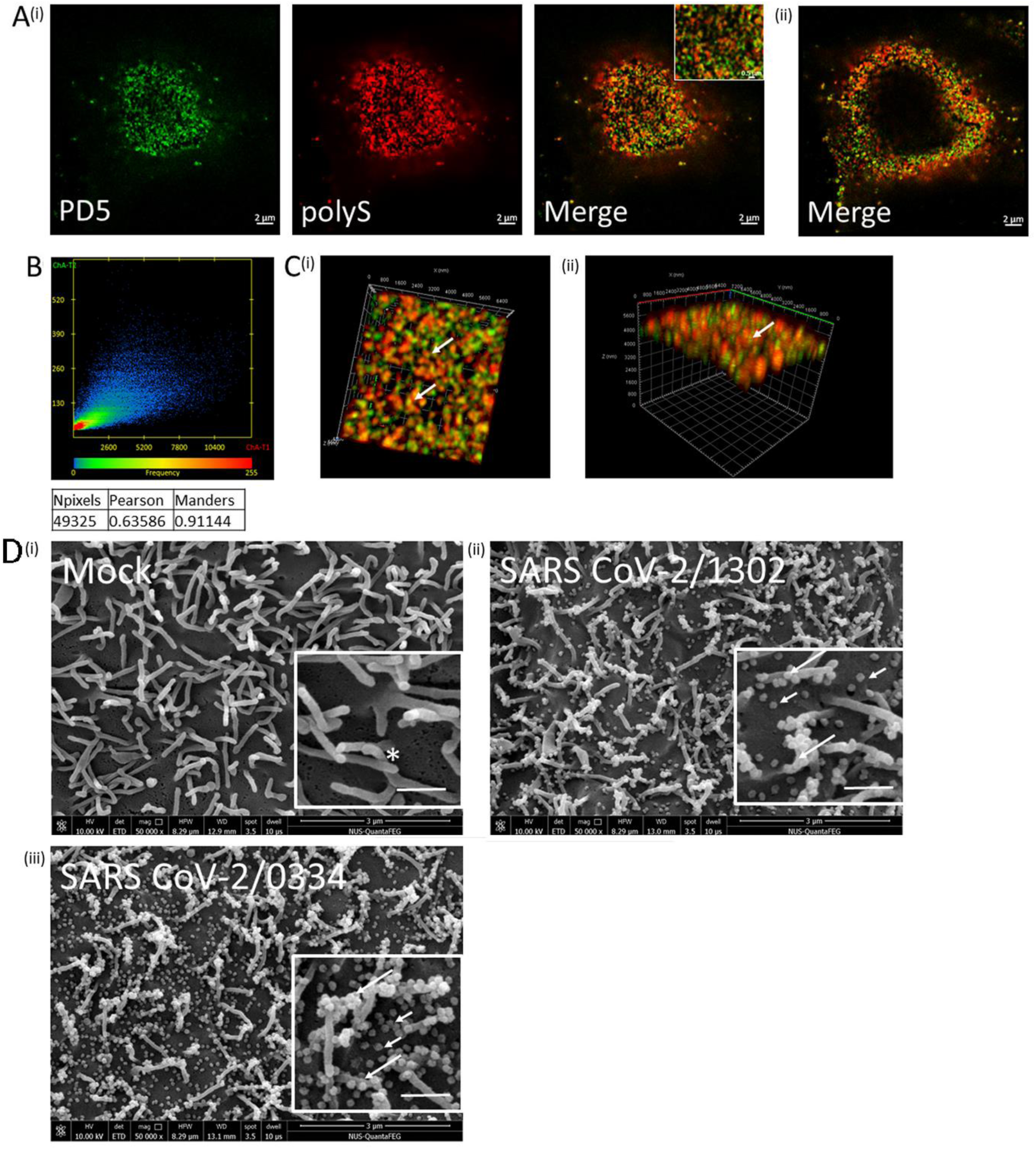
Distribution of PD5 and polyS staining on the surface of SARS-CoV-2 infected Vero E6 cells. Vero E6 cells were infected with SARS-CoV-2/0334 and at 18 hrs post-infection (hpi) the non-permeabilized cells were co-stained using PD5 and polyS. The co-stained cells were imaged using confocal microscopy and a series of images from the stained cell was obtained in the Z-plane. **(A)** Individual optical slices from the (i) cell top and (ii) at the cell periphery are shown. The individual channels and merged images are shown and an inset in (i) highlights the staining patterns at higher magnification. **(B)** The scatter plot and the values for the Pearson’s and Mander’s coefficients are shown from the 49,325 pixels sampled in the co-stained image. **(C)** The 3-dimensional reconstruction of the Z-stack series of images is shown. The reconstructed imaged is view from (i) above and (ii) at the side of the co-stained cell. The PD5 and polyS filamentous co-stained pattern on the cell surface is highlighted (white arrow). **(D)** (i) Mock-infected cells and cells infected with (ii) SARS-CoV-2/1302 and (iii) SARS-CoV-2/0334 at 18 hpi were imaged using scanning electron microscopy (50,000x magnification). In each case the associated inset is an enlargement (white bar =500nm). The individual virus particles on the cell surface (short arrow) and clusters of virus particles associated with cell microvilli (long arrows) are highlighted. The microvilli on the surface of mock-infected cells are also highlighted (*).

The surface topology of Vero E6 cells infected with SARS-CoV-2 isolates was further examined using scanning electron microscopy (SEM) to better understand the relevance of the different surface-staining patterns detected in the confocal microscope analysis described above. Cells were either mock-infected or infected with SARS-CoV-2/1302 and SARS-CoV-2/0334 isolates and at 18 hpi imaged using SEM (Fig. 5D). Numerous surface projections were observed on the surface of the mock-infected cells that was consistent with the presence of microvilli (Fig. 5D(i)), and these structures were also present on the surface of cells infected with SARS-CoV-2/1302 (Fig. 5D(ii)) and SARS-CoV-2/0334 (Fig. 5D(iii)). The SARS-CoV-2 particles are approximately 80-100 nm in diameter [34], and the numerous spherical particles of similar uniform dimensions that were detected on the surface of the virus-infected cells were consistent with the presence of individual SARS-CoV-2 particles. Approximately 70% of these virus particles were also associated with the microvilli projections, and we estimated that up to ten virus particles could be detected on many microvilli. Microvilli extending from individual cells to neighboring cells in the monolayer was a common occurrence on mock-infected cells (SFig 5A) indicating that under normal conditions these surface structures can make direct physical connections to neighboring cells. In this context, cells infected with either isolate showed virus particles on these microvilli extending from visibly infected cells to neighboring non-infected cells in the cell monolayer was noted (SFig. 5B-D), suggesting that these pre-existing surface projections may serve as a conduit to facilitate localized cell-to-cell transmission of SARS-CoV-2 in the cell monolayers. This is consistent with the recent interpretations of SARS-CoV-2 transmission in infected Vero cells monolayers [35, 36]. Although the significance of the association of SARS-CoV-2 particles with these cellular projections is uncertain, this explains the apparent filamentous PD5 and polyS co-staining pattern observed in the confocal microscopic analysis.

A comparison of the imaging data by light microscopy and SEM provided evidence that the surface staining correlated with the appearance of SARS-CoV-2 particles on the surface of infected cells. This indicated that surface-PD5 staining could be used to detect virus particles on the surface of SARS-CoV-2-infected cells. The surface staining of SC29 also provided evidence that the non-neutralizing antibodies can also bind to the virus particles on the surface of the infected cells. Although the RBD-binders recognize the S protein at the Golgi complex, the surface PD5 staining suggested that the antibodies also recognized the S protein in post-Golgi transport compartments in which the virus particles are transported to surface of infected cells.

### 4. Establishing the kinetics of cell-to-cell spread for SARS-CoV-2/1302 and SARS-CoV-2/0334 in the Vero E6 cell monolayers

The SEM analysis indicated that large numbers of SARS-CoV-2 particles were present on the surface of infected Vero E6 cells, suggesting that a high level of virus infectivity remained cell-associated at this time of infection. We estimated that on average greater than 200 virus particles/cell were detected on SARS-CoV-2/0334-infected cells, which is in the same order of magnitude for the estimated burst size for other coronaviruses [37]. In addition, we consistently noted lower numbers of virus particles on SARS-CoV-2/1302-infected cells compared with SARS-CoV-2/0334 virus-infected Vero E6 cells. Since the cells were exposed to similar levels of each virus isolate (moi of 0.1) and examined at the same time of infection, these differences suggested a slower rate of appearance of virus particles (i.e. virus particle assembly) in SARS-CoV-2/1302-infected Vero E6 cells and this was examined further. Vero E6 cell monolayers were infected with SARS-CoV-2/1302 or SARS-CoV-2/0334 using a moi of 0.001 (Fig.6A), 0.01 (Fig. 6B) and 0.1 (Fig. 6C), and at 18 hpi the cell monolayers were co-stained using PD5 and polyS and imaged by IF microscopy. At all moi values examined the two virus isolates exhibited an antibody staining pattern that was largely composed of clusters of stained cells. This was consistent with the localized cell-to-cell virus transmission, in which the progeny virus particles produced in the initial virus-infected cell infected the immediate surrounding cells in the monolayer. If high levels of infectious SARS-CoV-2 were shed into the tissue culture media covering the infected cells a more sporadic antibody labelling pattern would be expected that consisted of higher numbers of individual stained cells randomly distributed in the monolayer. It was noticeable that the cells infected with SARS-CoV-2/1302 consistently exhibited smaller infected cell clusters (10±1.5 cells per cluster; moi = 0.01) compared with the cells infected with the SARS-CoV-2/0334 isolate (25±2.1 cells per cluster; moi = 0.01) (Fig. 6D), which suggested reduced cell-to-cell transmission exhibited by SARS-CoV-2/1302. In this context cells infected with the SARS-CoV-2/0563 and stained using polyS also showed infected cell clusters that were of a similar size to those in SARS-CoV-2/0334-infected cells (SFig. 6A).

**Fig. 6.**
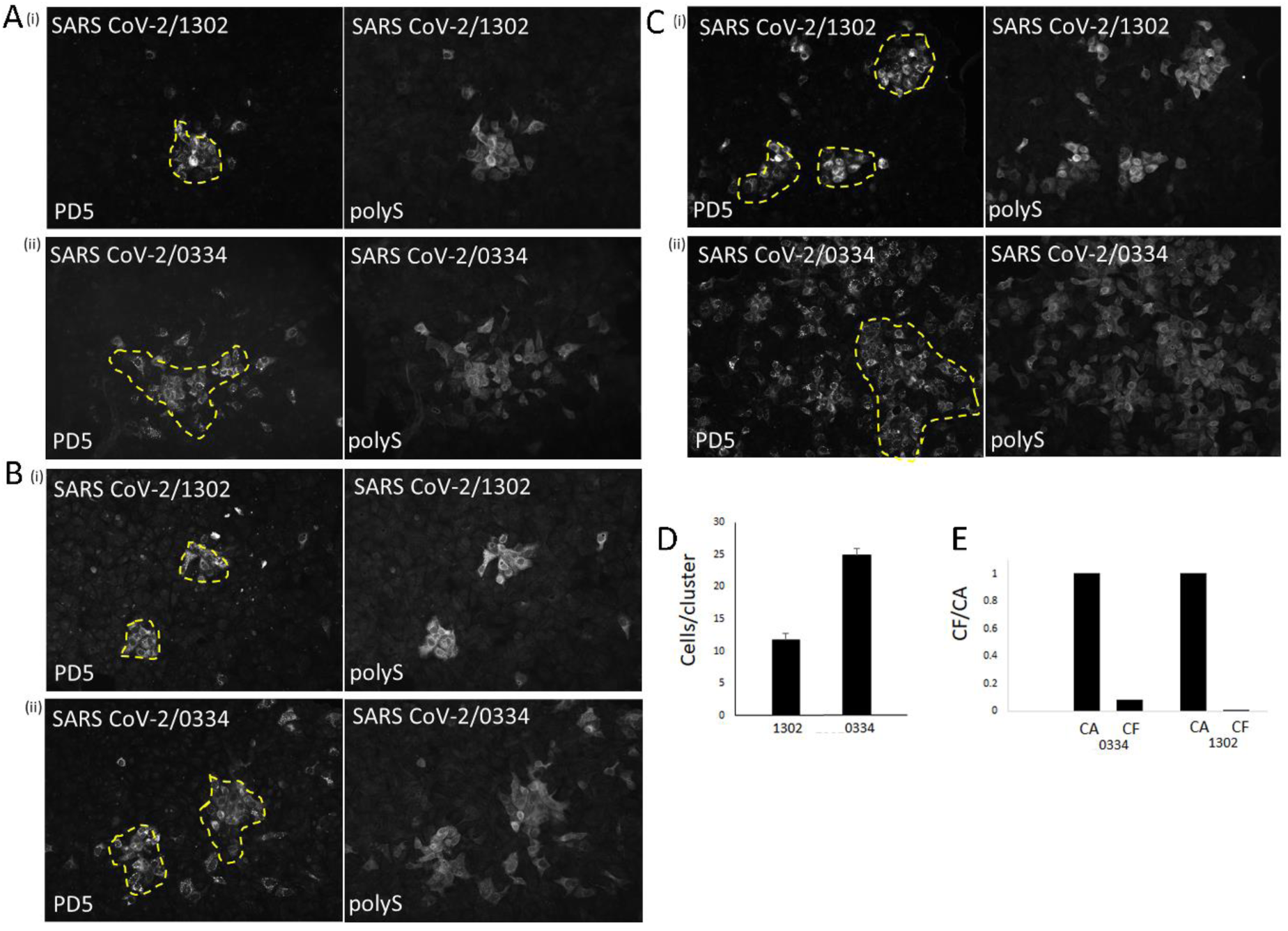
Comparison of Vero E6 cells infected with SARS-CoV-2/1302 and SARS-CoV-2/0334 at different multiplicity of infections. Vero E6 cells were infected with (i) SARS-CoV-2/1302 and (ii) SARS-CoV-2/0334 using a multiplicity of infection (moi) of **(A)** 0.001, **(B)** 0.01 and **(C)** 0.1, and at 18 hrs post-infection (hpi) the cells were co-stained using PD5 and polyS, and imaged using immunofluorescence microscopy (objective x20 magnification). In each condition the representative infected cell clusters are highlighted (broken yellow line). **(D)** The average numbers of infected cells per infected cell cluster in cells infected with SARS-CoV-2/1302 (1302) virus and SARS-CoV-2/0334 (0334) using a moi of 0.01 is shown. **(E)** The relative levels of cell-associated (CA) and cell-free (CF) virus infectivity recovered from cells infected with SARS-CoV-2/0334 (0334) and SARS-CoV-2/1302 (1302) using a moi of 0.01. In this analysis the results are presented as a fraction of the CA infectivity (which is set to a value of 1.0).

This clustered antibody staining was also observed on PD7, SC23 and SC29-stained SARS-CoV-2/1302 (SFig. 6B) and SARS-CoV-2/0334 (SFig. 6C) infected cells, indicating that the clustered staining pattern was not antibody-specific (i.e., due to PD5 and polyS staining). This interpretation of localized virus transmission of these SARS-CoV-2 isolates was consistent with recent observations of SARS-CoV-2 transmission reported by other laboratories [35]. The cell-to-cell transmission of these virus isolates was further supported by examining the relative level of virus-infectivity that was cell-free (tissue culture supernatant) and cell-associated (cell mass), and in cells infected with either isolate the cell-associated virus infectivity accounted for a higher proportion of the total infectivity (approximately 90% of the total recovered infectivity) (Fig. 6E). These data provide evidence for localized cell-to-cell transmission in the cell monolayer and suggest that SARS-CoV-2/0334 exhibited faster rate of transmission than the SARS-CoV-2/1302.

The kinetics of virus spread in the Vero cell monolayer was determined in Vero E6 cells infected with SARS-CoV-2/1302 (Fig. 7A) or SARS-CoV-2/0334 (Fig. 7B) using a moi of 0.01. At between 6 and 30 hpi the cells were co-stained with PD5 and polyS, and examined by IF microscopy. At 6 hpi we failed to detect significant levels of virus infectivity suggesting that this is at a time prior to significant levels of progeny virus production. At 18 hpi the appearance of infected cell clusters could be readily detected in cell monolayer infected with either isolate, which became progressively larger at 24 and 30 hpi. These rates of virus spread in the cell monolayers are consistent with previous reports that have examined the replication kinetics of SARS-CoV-2 in Vero cells [38]. However, at each time of infection the infected cell clusters in SARS-CoV-2/1302-infected cell monolayer were smaller than those in SARS-CoV-2/0334-infected cells, and was consistent with the slower spread of infection for SARS-CoV-2/1302. A similar analysis on Vero E6 cells infected with SARS-CoV-2/1302 (SFig. 7A) and SARS-CoV-2/0334 (SFig. 7B), and co-stained with PD7 and anti-NP again showed smaller infected cell clusters at all times of infection in SARS-CoV-2/1302 infected cell monolayers. Since all antibodies showed similar clustered staining patterns, the difference in rate of cell-to-cell spread between the two virus isolates was not antibody-specific e.g. due to differences in RBD recognition by PD5.

**Fig. 7.**
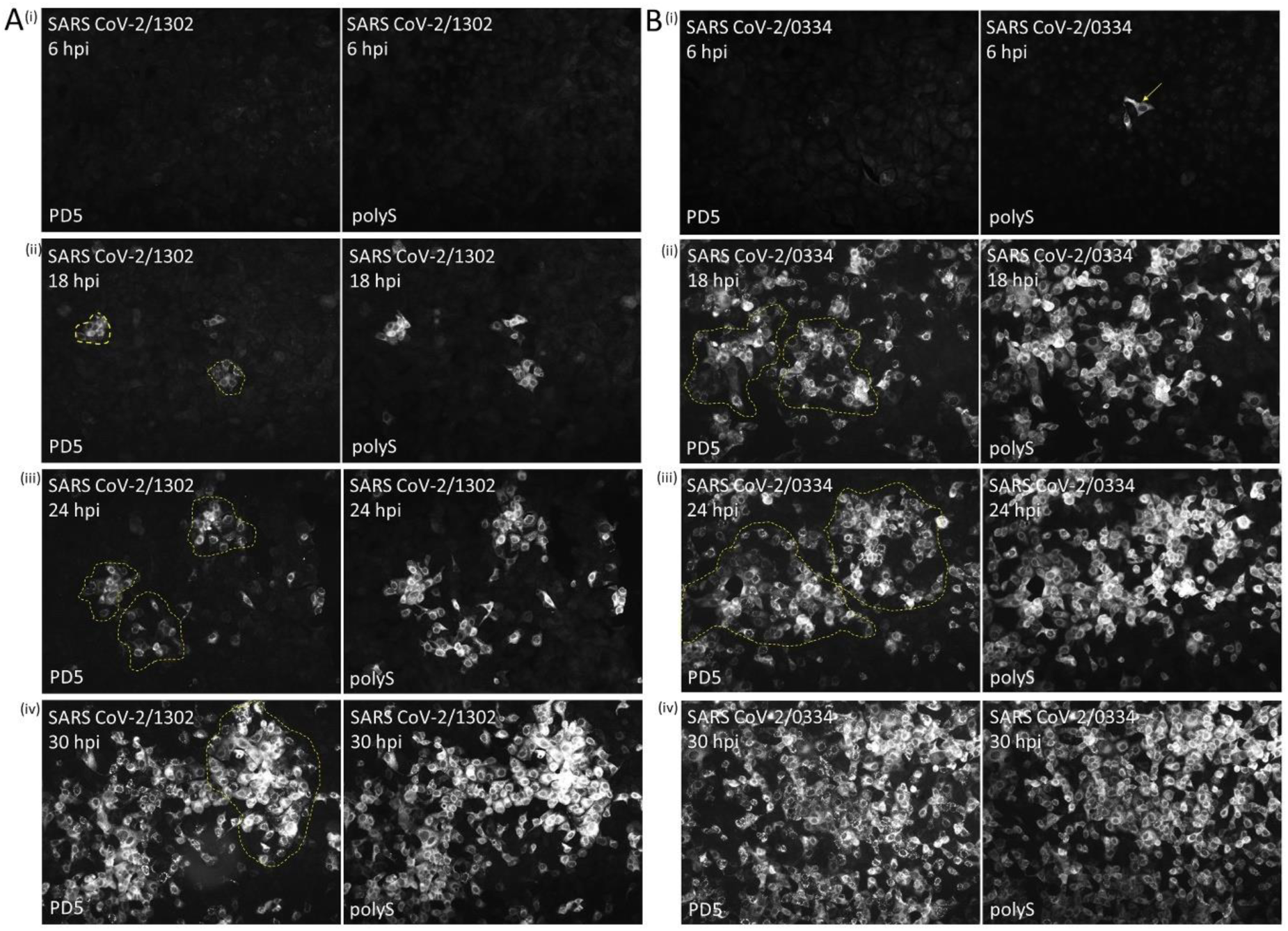
Temporal appearance of PD5 and anti-polyS staining on Vero E6 cells infected with the SARS-CoV-2/1302 and SARS-CoV-2/0334. Vero E6 cells were infected with **(A)** SARS-CoV-2/1302 virus and **(B)** SARS-CoV-2/0334 using a multiplicity of infection of 0.01 and at (i) 6 hrs post-infection (hpi), (ii) 18 hpi, (iii) 24 hpi and (iv) 30 hpi the cells were co-stained with PD5 and polyS. The stained cells were imaged using immunofluorescence microscopy (objective x20 magnification). A sporadic infected cell at 6 hpi is highlighted (white arrow), and in both **(A)** and **(B)** clusters of infected cells are highlighted (broken yellow line).

Our data is consistent with localized cell-to-cell spread in the cell monolayer that correlated with high levels of cell-associated virus. However the reason for the high level of cell-associated virus at this time of infection is currently uncertain and will require further investigation. In this context, high resolution structures of the S protein on isolated SARS-CoV-2 particles have demonstrated that low levels of the S protein trimer disassociate leaving the S2 domain radiating from the virus envelope in an extended post-fusion conformation [39]. It is possible that the freely exposed fusion peptide at the N-terminus of the S2 protein may insert into the plasma membrane on infected cells and serve as an alternative transmembrane domain [40]. In such a scenario, this would tether the virus particles to the surface of infected cells and facilitate the localized cell-to cell spread of infection that we observe. The reason for the different rate of spread in the Vero cell monolayer is still uncertain. We failed to detect a significant difference in the time of appearance of the antibody stained cells infected with either isolate in these cells. This suggests that the differences in virus spread may be due to differences between the isolates at the later stages of the virus replication, however the mechanism behind this phenomenon will require further investigation.

### 5. N-linked glycan maturation or furin cleavage of the S protein is not required for recognition of the RBD by PD5 in SARS-CoV-2 infected Vero E6 cells

Furin is enriched at the Golgi complex [41], and post-translational cleavage of the S0 protein into the S1 and S2 domains at the S1/S2 site would be expected to occur as it is trafficked through the Golgi complex. Furthermore, processing of the N-linked glycan from high-mannose cores to terminally differentiated complex glycans also occurs in the Golgi complex. The S protein contains several potential N-linked glycosylation sites, including two sites within the RBD. We therefore examined if maturation of the associated N-linked glycans and furin cleavage of the S protein was required for the recognition of the RBD by PD5.

Inhibition of the processing of S protein N-linked glycans into complex glycans was performed using the Golgi mannosidase-1 inhibitor deoxymannojirmycin (DMJ) (SFig. 8). We have previously used DMJ to examine glycan heterogeneity in of the RSV F protein in Vero cells [42, 43]. The recombinant S protein expressed in non-treated cells and in DMJ-treated cells was examined by immunoblotting using polyS (SFig. 8A(i)). A band shift in the migration of the S2 subunit (increased electrophoretic migration) in DMJ-treated cells was consistent with the presence of high mannose cores attached to the S protein [42, 43]. This was confirmed by examining sensitivity of the recombinant S protein associated glycans to digestion with the enzyme endoH (SFig. 8A(ii)). This enzyme removes high mannose cores from glycoproteins but is unable to remove complex N-linked glycans, and is thus an established reagent to examine glycan maturation in glycoproteins. In non-treated cells the S protein exhibited partial sensitivity to endoH cleavage suggesting that individual S protein polypeptide chains contained a mixture of simple and complex glycans as described previously ([44–46]. In the DMJ-treated cells the S2 subunit exhibited total sensitivity to endoH treatment as indicated by the increased migration of the S2 protein, and which was consisted with the S protein in DMJ-treated Vero cells exhibiting only simple high mannose cores. In cells expressing either the recombinant S protein (SFig. 8B) or in cells infected with SARS-CoV-2/0334 (SFig. 8C and D) the S protein exhibited similar polyS and PD5 staining in non-treated and DMJ-treated cells. This indicated that PD5 recognition was not dependent on the conversion of the high-mannose N-linked glycans into complex N-linked glycans, although it is acknowledged that other types of modification such as O-linked glycosylation may play a role in facilitating PD5 binding.

We have previously used the furin inhibitor decanoyl-RVKR-cmk (RVKR) to examine furin cleavage of the respiratory syncytial virus fusion protein in Vero E6 cells [47]. Vero E6 cells expressing the recombinant S protein was used to determine the effective concentration of decanoyl-RVKR-cmk to inhibit S protein cleavage. At 4 hr post-transfection (hpt) the cells expressing the recombinant S protein were either non-treated or treated with 10, 20 or 40 µM RVKR, and at 16 hpt the post-translational cleavage of the S protein was examined by immunoblotting cell lysates with polyS (Fig. 8A). In non-treated cells both S0 and S2 was detected indicating post-translation cleavage of the S protein. Reduced levels of S2 protein were detected as the RVKR concentration was increased from 10 to 40 µM, and at 40 µM no residual S protein cleavage was detected. Under our experimental conditions we failed to detect any reduction in the S0 protein levels after drug-treatment, which suggested that under these experimental conditions there was minimal drug toxicity and was consistent with the LDH cytotoxicity assay which indicated 95% cell viability (Sugrue unpublished observations). In a parallel analysis, non-treated and RVKR-treated cells expressing the recombinant S protein were also co-stained with polyS and PD5, and imaged using IF microscopy (Fig. 8B(i) and (ii)). Large antibody-co-stained cell clusters in non-treated cells indicated the formation of the multinucleated cell clusters, which suggested that expression of the recombinant S protein alone may be sufficient to induce membrane-fusion in Vero E6 cells and is consistent with recent reports [48, 49]. In A549 cells expressing the recombinant S protein we failed to detect the presence of these multinucleated cells, although PD5 and polyS staining was observed (Lim and Sugrue, unpublished observations). The A549 cells do not express the ACE2 protein, and we presume that in Vero cells the recombinant S protein engages with the ACE2 receptor and is able to induce receptor-mediated membrane fusion. In this context the SARS-CoV-2 S protein is expected to bind to the ACE2 protein of several different animal species, including primates from which the Vero cell line is derived [50–52]. Although furin cleavage of the S protein appears not an absolute requirement for mediating membrane fusion [48], we failed to detect the presence of multinucleated cells in the PD5 and polyS co-stained cells RVKR-treated cells. The change in staining pattern following RVKR-treatment suggested a correlation between S protein cleavage and membrane fusion. However, the PD5 staining in both non-treated and RVKR-treated cells indicated that furin cleavage of the recombinant S protein was not required for RBD recognition by PD5.

**Fig. 8.**
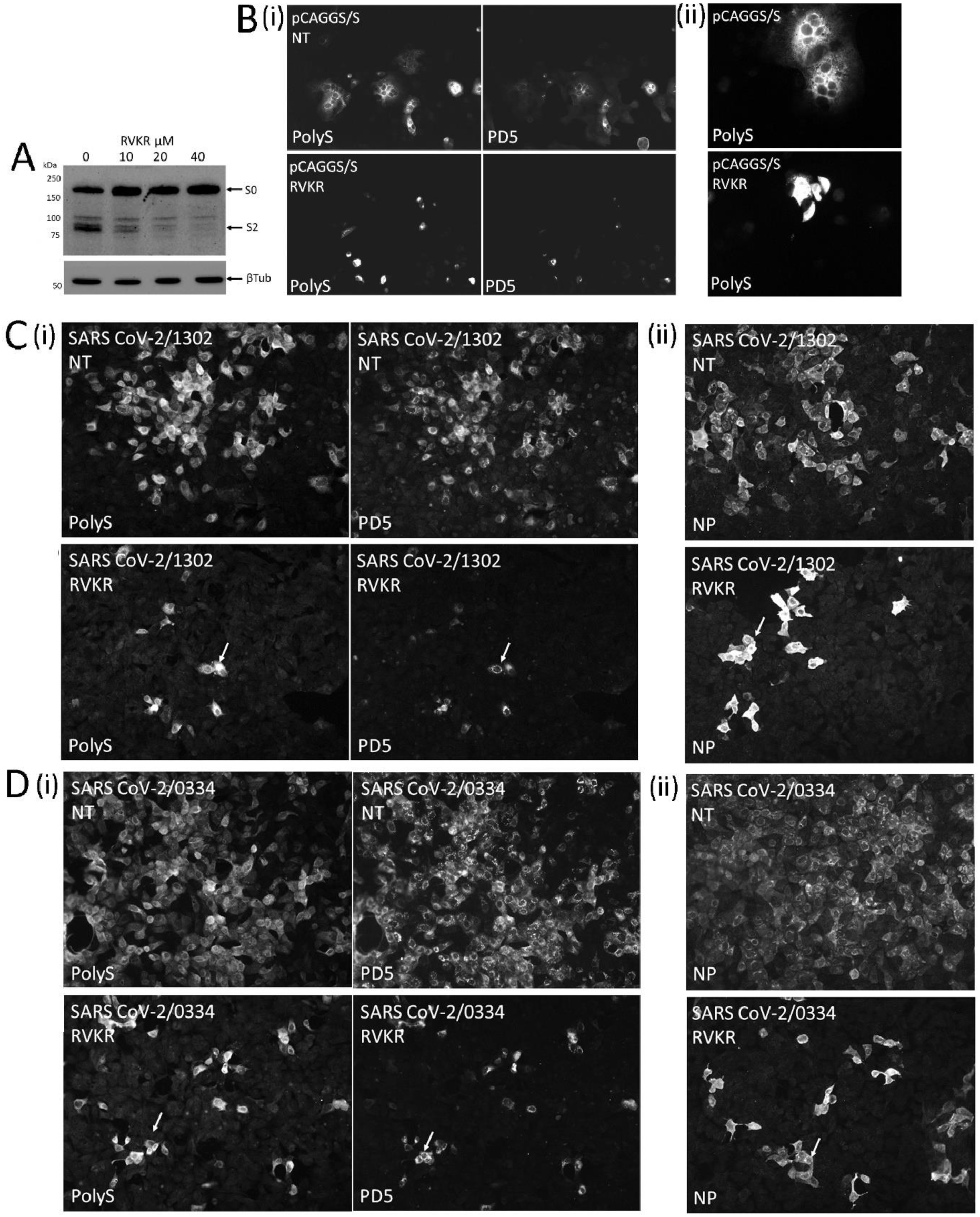
Furin cleavage of the S protein is not required for RBD recognition by PD5. **(A)** Vero E6 cells expressing the recombinant S protein were treated with 0, 10, 20 and 40μM decanoyl-RVKR-cmk (RVKR) at 4 hrs post-transfection (hpt) and at 16 hpt, the cell lysates were prepared and immunoblotted with polyS. The full-length S protein (S0) and S2 subunit are indicated. Tubulin (βTub) is the loading control. **(B)** (i) Non-treated (NT) and 40μM RVKR-treated cells expressing the recombinant S protein were stained with PD5 and polyS, and imaged using immunofluorescence (IF) microscopy (objective x20 magnification). (ii) Imaging of polyS-stained NT and RVKR-treated cells expressing the recombinant S protein at 16 hpt (objective x40 magnification). Cells were infected with **(C)** SARS-CoV-2/1302 and **(D)** SARS-CoV-2/0334 using a multiplicity of infection of 0.01 in the absence (NT) and presence of 40μM RVKR. At 18 hpi, the cells were co-stained using (i) PD5 and polyS, and (ii) anti-NP and PD5, and imaged using IF microscopy (objective x20 magnification). The reduced size of the infected cell clusters in RVKR-treated cells is highlighted (white arrow).

The effect of RVKR-treatment on PD5 recognition in Vero E6 cells infected with SARS-CoV-2/1302 (Fig 8C) and SARS-CoV-2/0334 (Fig 8D) was also examined. Virus-infected Vero E6 cells were either non-treated or treated with 40 µM RVKR at 4 hpi, and at 18 hpi the cells were co-stained with polyS and PD5 (Fig. 8C(i) and D(i)) and with anti-NP (Fig. 8C(ii) and D(ii)). The RVKR-treatment was started at 4 hr after infection to ensure no interference by the drug in establishing the initial SARS-CoV-2 infection, but was added at a time that was prior to the formation of the infected cell clusters. In the non-treated cells widespread antibody staining within the cell monolayers indicated the spread of virus infection in the cell monolayer infected with both isolates. In RVKR-treated cells the antibody staining with either isolate was restricted to individual brightly stained infected cells and smaller infected cell clusters (approximately 2-3 infected cells per cluster). However, RVKR-treatment did not prevent PD5 recognition of the S protein and indicated that in virus-infected cells furin-cleavage of the S protein was not a requirement for RBD binding by PD5. Although the S protein of SARS-CoV-2/1302 has a modified furin cleavage site, proteolytic cleavage of the S protein by furin was still required for its transmission. These data underpin the importance of the furin cleavage of the S protein in mediating virus transmission in animal models of infection [53, 54].

In a final analysis we examined if furin cleavage of the S protein was required for the recognition of the RBD by PD5 on the cell surface of Vero E6 cells infected with SARS-CoV-2/1302 (Fig. 9A(i)) and SARS-CoV-2/0334 (Fig. 9A(ii)). While PD5 staining on non-treated cells was detected, a reduced level of PD5 surface staining on drug treated cells infected with either virus isolate was noted. Quantification of the PD5 staining intensity on SARS-CoV-2/0334-infected cells indicated an approximate 80% reduction in PD5 staining in RVKR-treated cells (Fig. 9A (iii) and (iv)). Individual representative non-treated and RVKR-treated non-permeabilized and permeabilized virus-infected cells that were co-stained with PD5 and polyS was also examined using confocal microscopy and a series of images were recorded in the Z-plane (SFig. 9). An individual representative image slice was extracted from the image series to highlight the differences in the surface-staining intensity (Fig.9B(i) and cytoplasmic-staining intensity (Fig. 9B(ii)) under each experimental condition. This showed a similar level of cytoplasmic PD5 staining in non-treated or RVKR-treated infected cells, but a reduction in the PD5 (and polyS) surface staining in RVKR-treated cells.

**Fig. 9.**
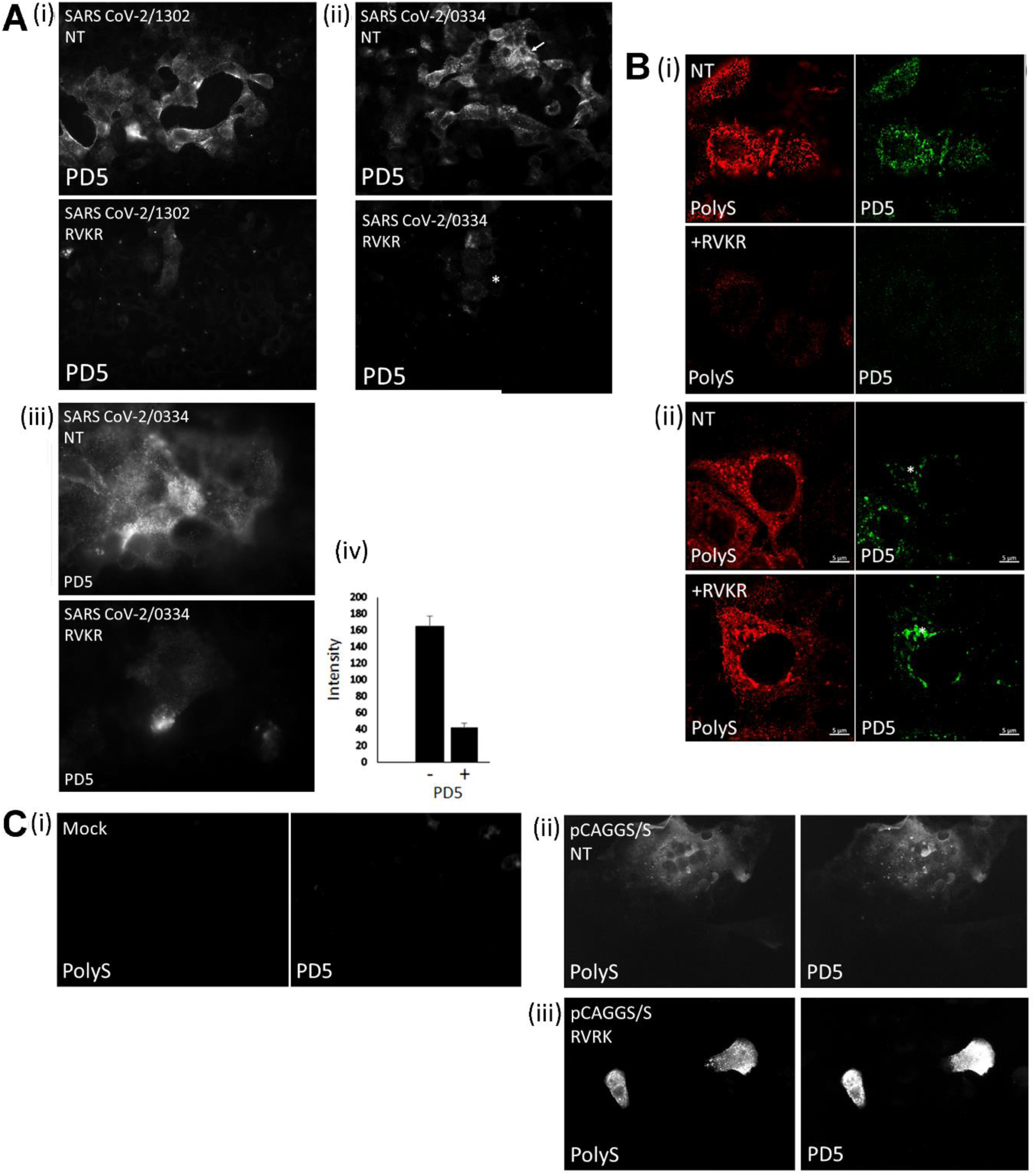
Furin cleavage of the S protein is required for surface display of the PD5 epitope in virus-infected Vero E6 cells. **(A)** Vero E6 cells were infected with (i) SARS-CoV-2/1302 and (ii) SARS-CoV-2/0334 and either non-treated (NT) or treated with 40μM decanoyl-RVKR-cmk (RVKR). At 18 hrs post-infection (hpi), the cells were non-permeabilized and stained with PD5 and imaged by immunofluorescence (IF) microscopy (objective x20 magnification). The reduced staining in RVKR-treated cells is highlighted (*). (iii) Non-treated (NT) and RVKR-treated cells infected with SARS-CoV-2/0334 were non-permeabilized and stained with PD5 and imaged by IF microscopy. (objective x100 magnification; oil immersion). (iv). The average image intensity of non-treated (-) and RVKR-treated (+) SARS-CoV-2/0334- infected cells stained with PD5 is shown. n=50 in each case. **(B)** The non-treated (NT) and RVKR-treated (+RVKR) SARS-CoV-2/0334-infected cells were (i) non-permeabilized and (ii) permeabilized, and co-stained with PD5 and polyS. An individual image taken from a Z-stack series of images of representative cells is shown, and the individual channel and merged images are presented. The punctate cytoplasmic PD5 staining pattern (*) is highlighted. **(C)** Cells were (i) mock-transfected or transfected with pCAGGS/S and either (ii) non-treated (NT) or (iii) RVKR-treated (RVKR) or (iii). The non-permeabilized cells were co-stained with PD5 and polyS, and imaged by IF microscopy (objective x40 magnification).

To determine if furin cleavage of the S protein was an essential requirement for its trafficking to the plasma membrane, non-permeabilized mock-transfected cells (Fig. 9A(i)) and cells expressing the recombinant S protein in non-treated (Fig. 9C(ii)) and RVKR-treated (Fig. 9C(iii)) cells were co-stained using polyS and PD5, and imaged using IF microscopy. Surface staining was detected on both non-treated and RVKR-treated cells using both antibodies, indicating that drug-treatment did not prevent surface expression of recombinant expressed S protein, and nor did it prevent surface-display of the RBD. The surface staining on non-treated cells appeared as numerous large antibody-stained clusters that was similar in appearance to the antibody-stained cells clusters detected in permeabilized cells, and was consistent with S protein-mediated membrane fusion. In contrast, on RVKR-treated cells the surface staining consisted of both single stained cells and much smaller stained cell clusters which was consistent with a reduced levels of membrane fusion. The RVKR-treated cells expressing the S protein also exhibited an apparent increased surface-staining intensity when compared with that in non-treated cells. Nevertheless, the recombinant S protein expression data indicated that furin cleavage of the S protein was not an inherent requirement for its trafficking to the plasma membrane.

It has been suggested that the S protein can be trafficked to the plasma membrane in virus infected cell without being incorporated into virus particles, and that this has been proposed to induce syncytia to facilitate virus transmission. However, our data suggests that cleavage of the S protein was required for its efficient surface expression in virus-infected cells which correlated with impaired virus cell-to-cell transmission in RVKR-treated infected cells. This may be related to other specific aspect of the virus replication cycle that prevents trafficking of the uncleaved S protein to the cell surface. For example, the S protein interacts with the M protein during virus particle formation [55], and in this context the uncleaved S protein may lead to the formation of aberrant virus particles that cannot form at the cell surface.

## Conclusion

The PD5, PD7 and SC23 that recognise the RBD of the S protein in virus-infected cells appear to recognise only a subpopulation of the S protein that has a distinct conformation in which the RBD is accessible to antibody binding. Although much of our characterisation has involved the use of PD5, the similarities in staining pattern between PD5 and the other RBD-binders suggested that PD5 was representative of these antibodies. The RBD-binders recognise the S protein at specific locations within the cell, which includes the Golgi complex and the virus particles that form at the cell surface. We failed to detect PD5 co-staining with antibodies that recognise the ER compartment (Sugrue unpublished observations) indicating that RBD recognition occurs in the Golgi complex and at post-Golgi cell compartments. This further suggests that during the transit of the S protein through the Golgi complex a conformational change in the S protein may occur that enables PD5 binding. In contrast the recognition of the S protein by PD4 and SC29 was not dependant on RBD accessibility, and these antibodies exhibit a widespread and diffuse staining pattern in virus-infected cells. Although PD4 and SC29 do not bind to the RBD, an additional punctate staining pattern that resembles that in the RBD-binders was also apparent, suggesting that they also recognise the form of the S protein that is recognised by the RBD-binders. We do not know which domain PD4 and SC29 bind in the S protein trimer, but our data indicates that they are able to recognise multiple different forms of the S protein in a manner similar to polyS antibody. There are currently several high-resolution structures that have been described that have used the soluble trimeric S protein ectodomain of both the uncleaved (S0) and the furin cleaved (S1/S2) forms of the S protein [22, 23]. In these structures it has been reported that one of the RBD in the S protein trimer can be elevated above the S protein trimer into an open conformation, and that the transformation of the RBD into the open form maybe associated with increased antibody accessibility [56]. These studies have suggested that the uncleaved and cleaved forms of the S protein with the RBD were preferentially in the closed and open conformation respectively. Interestingly, structural studies on the full-length S protein trimer complex suggest that in the uncleaved S protein the RBD also exists in the closed conformation [39, 57, 58], and this is likely to better reflect the conformation of the S protein in virus-infected cells. However, electron tomography on purified virus particles has highlighted the complexity of this issue, since these reports suggest the existence of multiple S protein conformations, including S protein trimers in a post-fusion conformation consisting only the S2 domain [39]. Although our data suggests that the RBD is first detected in the Golgi complex it is currently unclear if in the distinct S protein forms that are recognized by the PD5, PD7 and SC23 is related to the orientation of the RBD. It is possible that these antibodies recognise an alternative conformational change in the S protein during virus particle assembly that is not related to the elevation of the RBD. We can speculate that since the RBD-binders recognise conformational specific epitopes within the RBD these antibodies are able to detect formation of the correctly folded RBD prior to or during virus particle assembly. Future work will focus on identifying the precise binding sites of these antibodies in the S protein trimer to better understand the conformational changes in the S protein that is recognised by the RBD-binders in virus-infected cells.

## Limitations of this study

The work described in this study primarily relies on the use of Vero E6 cells, which is an established and accepted highly permissive cell system used to propagate SARS-CoV-2. These cells lead to the production of easily definable SARS-CoV2 particles and are useful to examine the mechanics of SARS-CoV2 infection. In this context important post-translational modification of the S protein such as furin cleavage and N-linked glycosylation also occurs in these cells. However, the physiology of these non-human cells are likely to exhibit differences when compared with the corresponding cells that are naturally infected in the airway of the human host. This may influence both the virus replication characteristics and processing of the individual virus proteins. Future work will characterise these antibodies in cells that are more representative of the human airway (e.g., nasal epithelial cells), to obtain a complete and more physiological picture of SARS-CoV-2 S protein antibody recognition.

We have used low passaged clinical virus isolates that were isolated using tissue culture during the early stages of the pandemic in Singapore. The focus of this study was on the S protein, and in this context the S protein sequence of the individual virus isolates used in this study were genetically characterised. Although we did observe some differences in the S protein sequences that were restricted to the vicinity of the furin cleavage site, the virus isolates showed high levels of sequence identity with the corresponding S protein sequence of SARS CoV-2/WIV04, including the sequence of the RBD. Understanding the complex biology of these virus isolates was not within the scope of the current study, but the differences that we observed in the localised virus transmission of these virus isolates in tissue culture suggest subtle differences in their replication characteristics. While it is unclear how genetic variation in other virus genes would directly influence the recognition of the S protein by the antibodies used in this study, future work on the complete genetic characterisation of these viruses may help to explain these differences in their biological properties that we observe.

Lastly, we have selected a panel of antibodies based in their binding properties to the S protein RBD. Although binding to the RBD was established for PD5, PD7 and SC23, we have not yet mapped their antibody binding sites on the RBD. This information should provide a much better understanding of the conformational changes in the S protein in virus-infected cells that we describe, and we will use established methodology to map their antibody binding sites on the S protein RBD in future work.

## Contribution from authors

Conceptualisation, RJS, CEZC, BHT; Methodology, APCL, SLKS, DHC, SKKW, CGN, JHL, VZYL, PSW, KML, VZYL, SKL, YCL; Data analysis, CEZC, CGN, BHT, RJS; Data curation, BHT, RJS; writing, reviewing, editing BHT, RJS; project administration, BHT, CEZC.

## Acknowledgements

The work was supported by DSO National Laboratories, and Nanyang Technological University. We acknowledged Sok-Kiang Lau, Jin-Phang Loh, Wee-Hong Koh, Hwee-Teng Low, Jasper Chin-Wen Liaw, Janet Seok-Wei Chew, Xiao-Fang Lim, Elizabeth Ai-Sim Lim Tan Jie Ling, and Sian-Foong Ling for technical assistance and the diagnostics performed on the samples and the Facility Team (Loh Siang Gary, Mok-Wei Heng, Chang-Lin Chang, and Lay-Tin Aw) for their assistance.

## SFigures

**SFig. 1.**
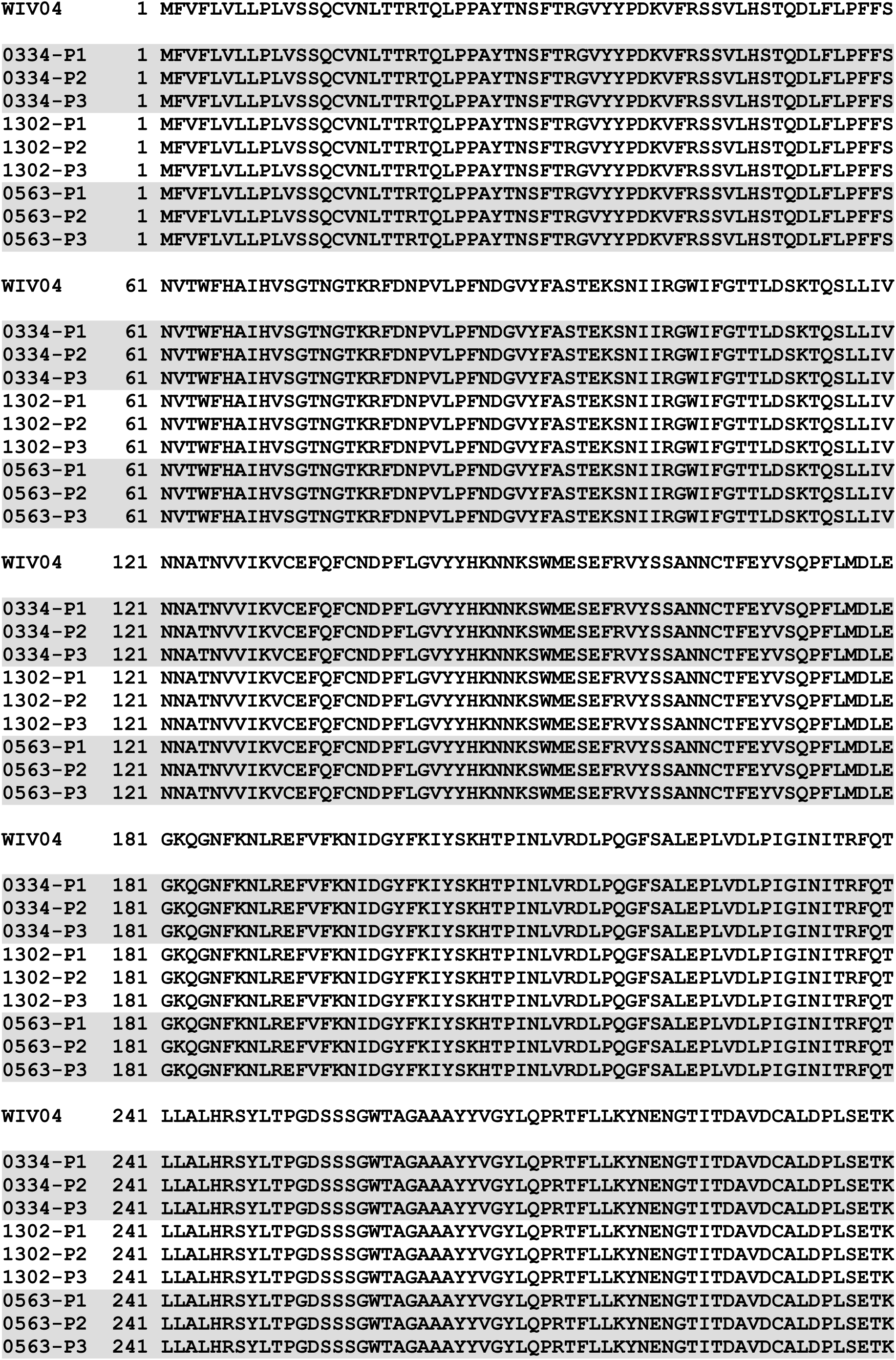
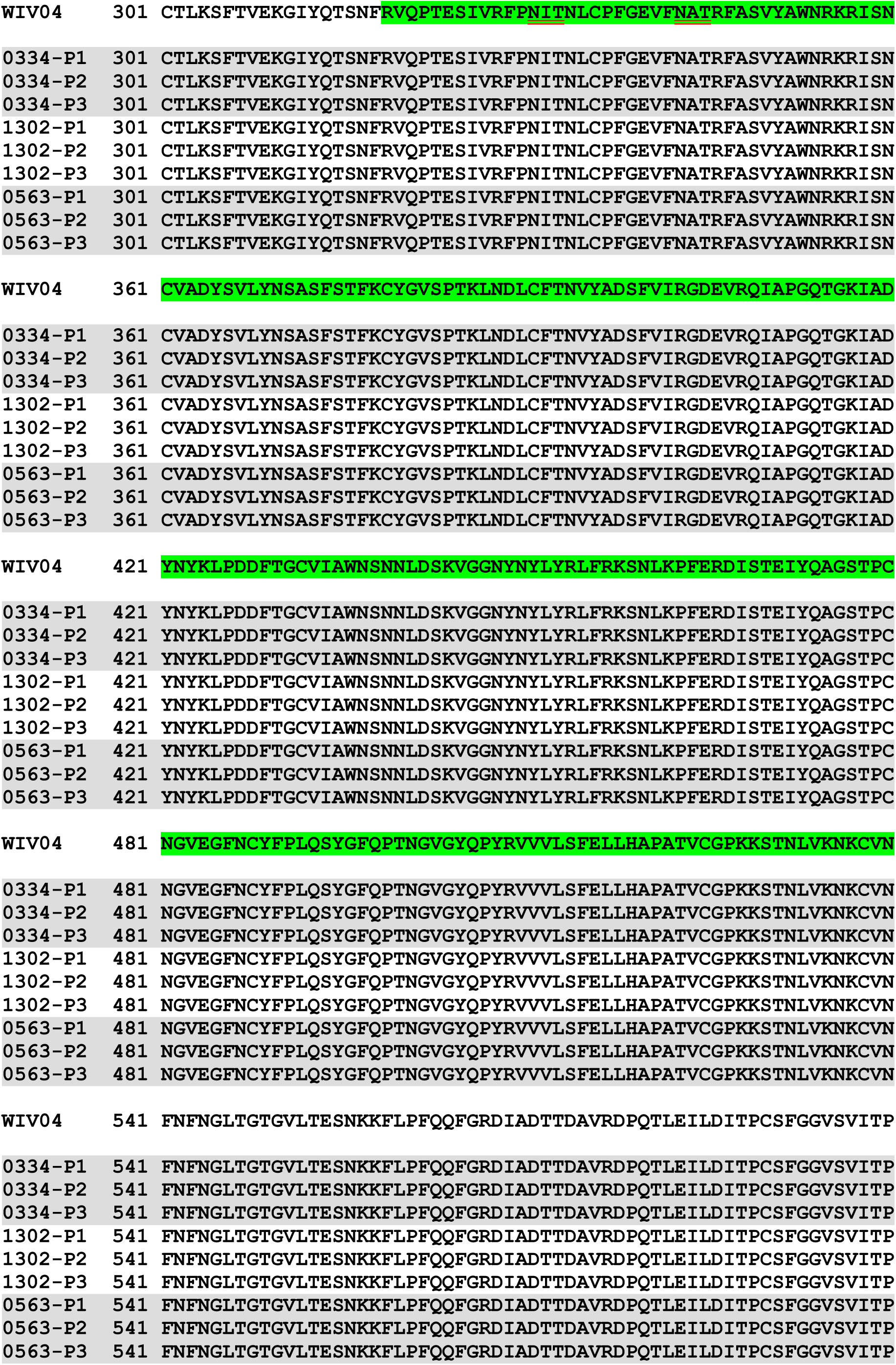
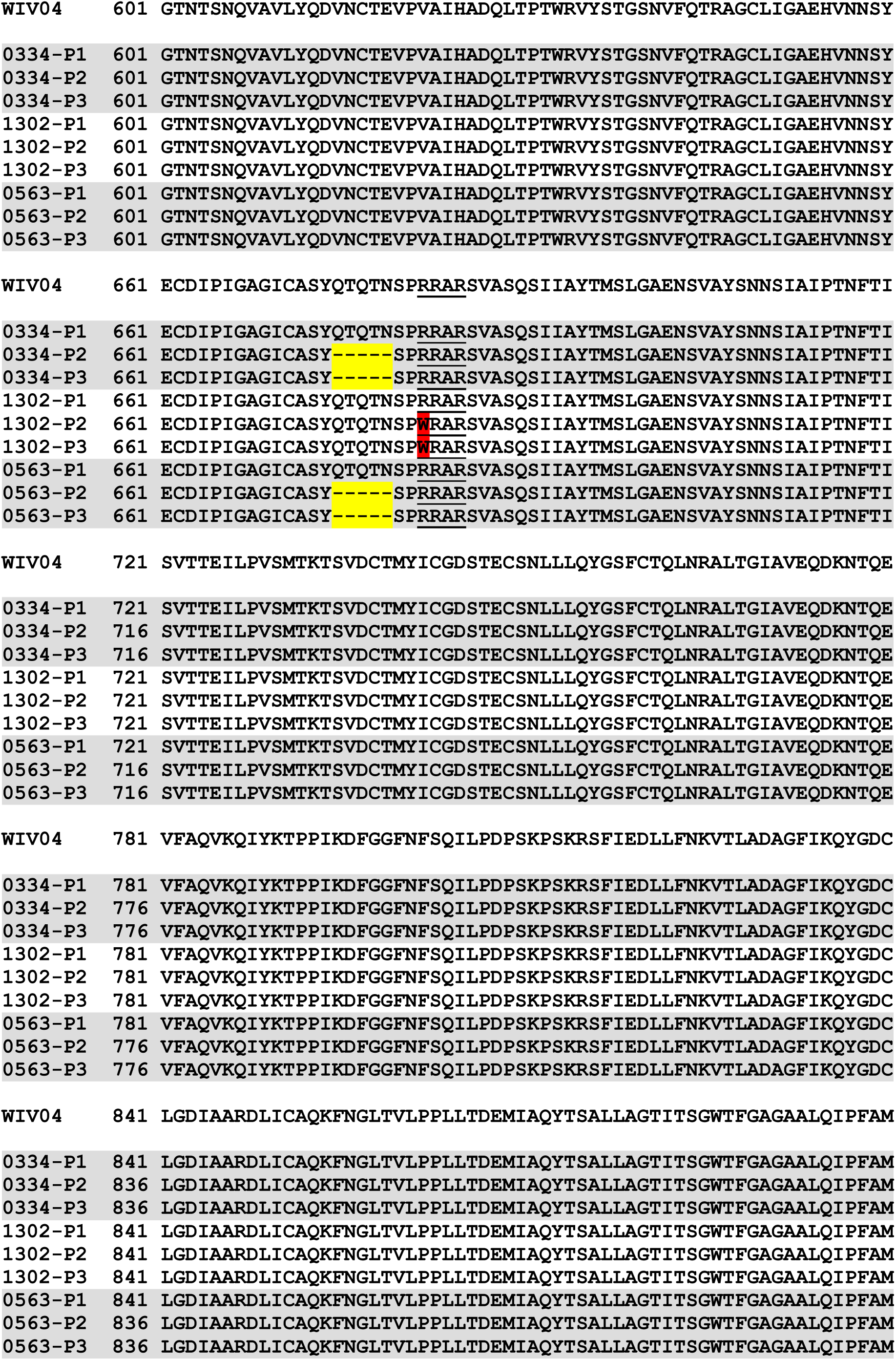
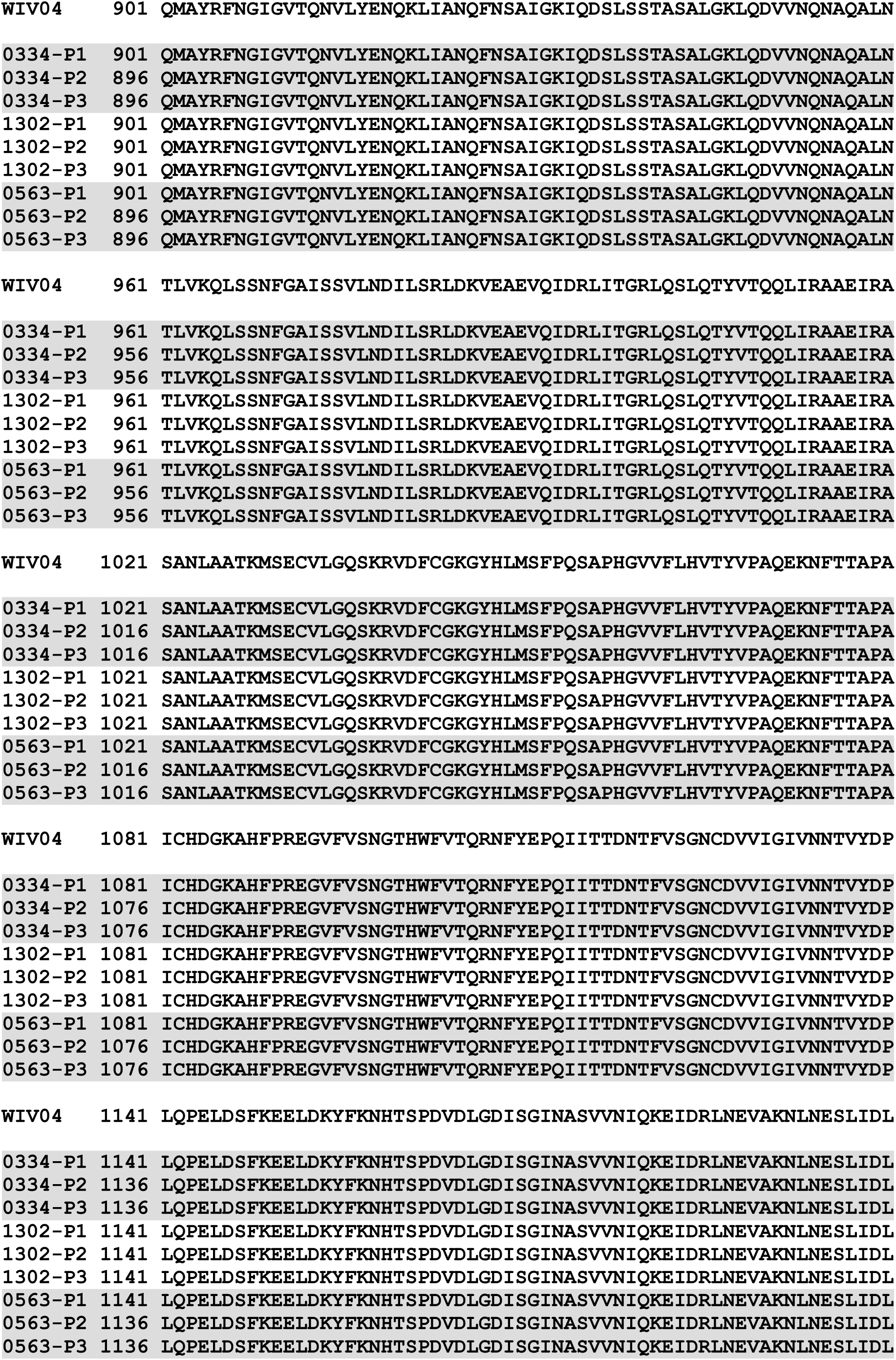
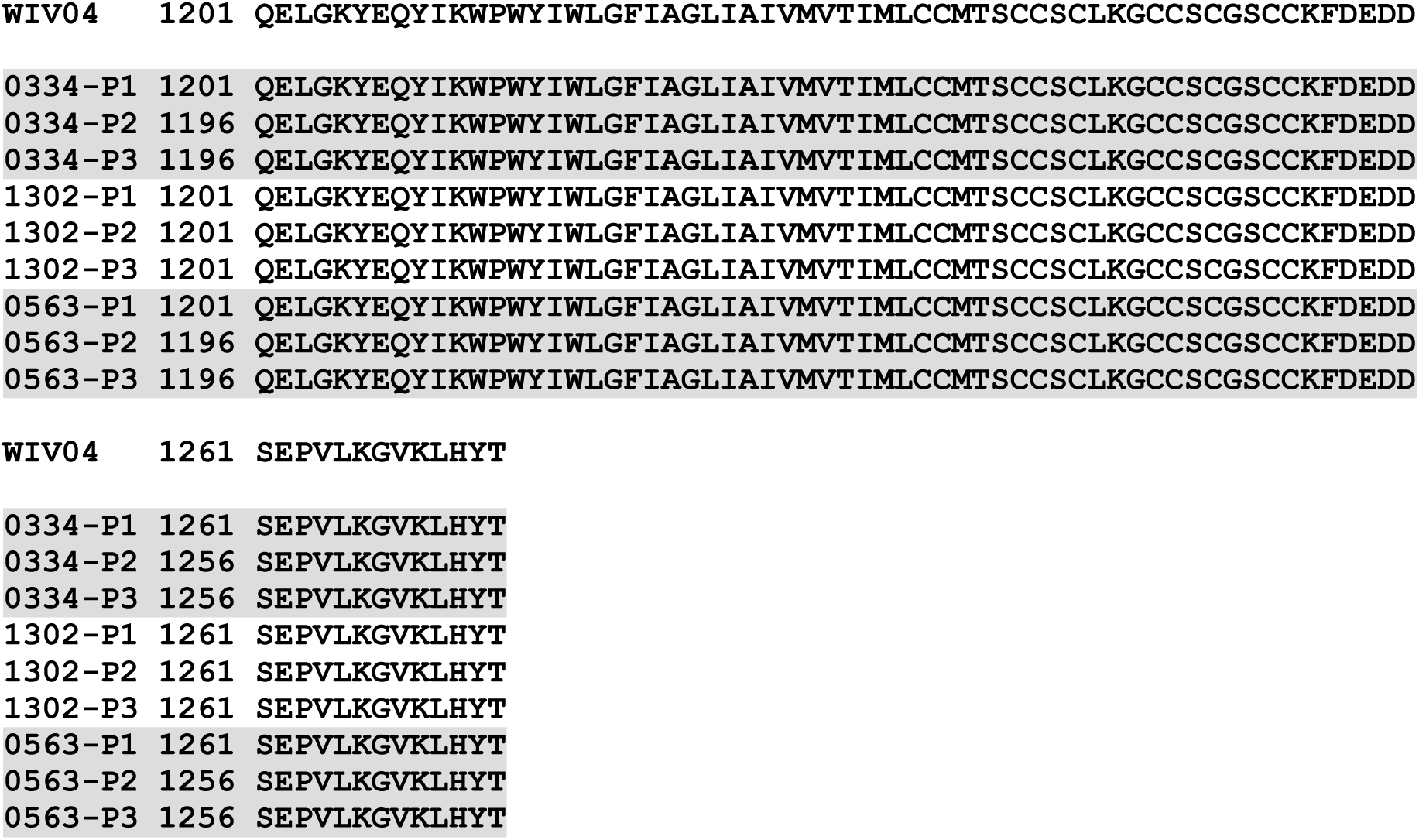
Amino acid sequence of the S protein of SARS-CoV-2/0334, SARS-CoV-2/0563 and SARS-CoV-2/1302. The S protein sequence alignment is shown for SARS-CoV-2 (WIV04), SARS-CoV-2/0334, SARS-CoV-2/0563, and SARS-CoV-2/1302. The receptor binding domain (RBD) at amino acid positions 319-540 (green highlight), deleted sequence at amino acid position 675-679 (dashed line; yellow highlight), the two potential N-linked glycosylation sites in the RBD (double underline red lines), and the mutation at position R682W (red highlight) are indicated.

**SFig 2.**
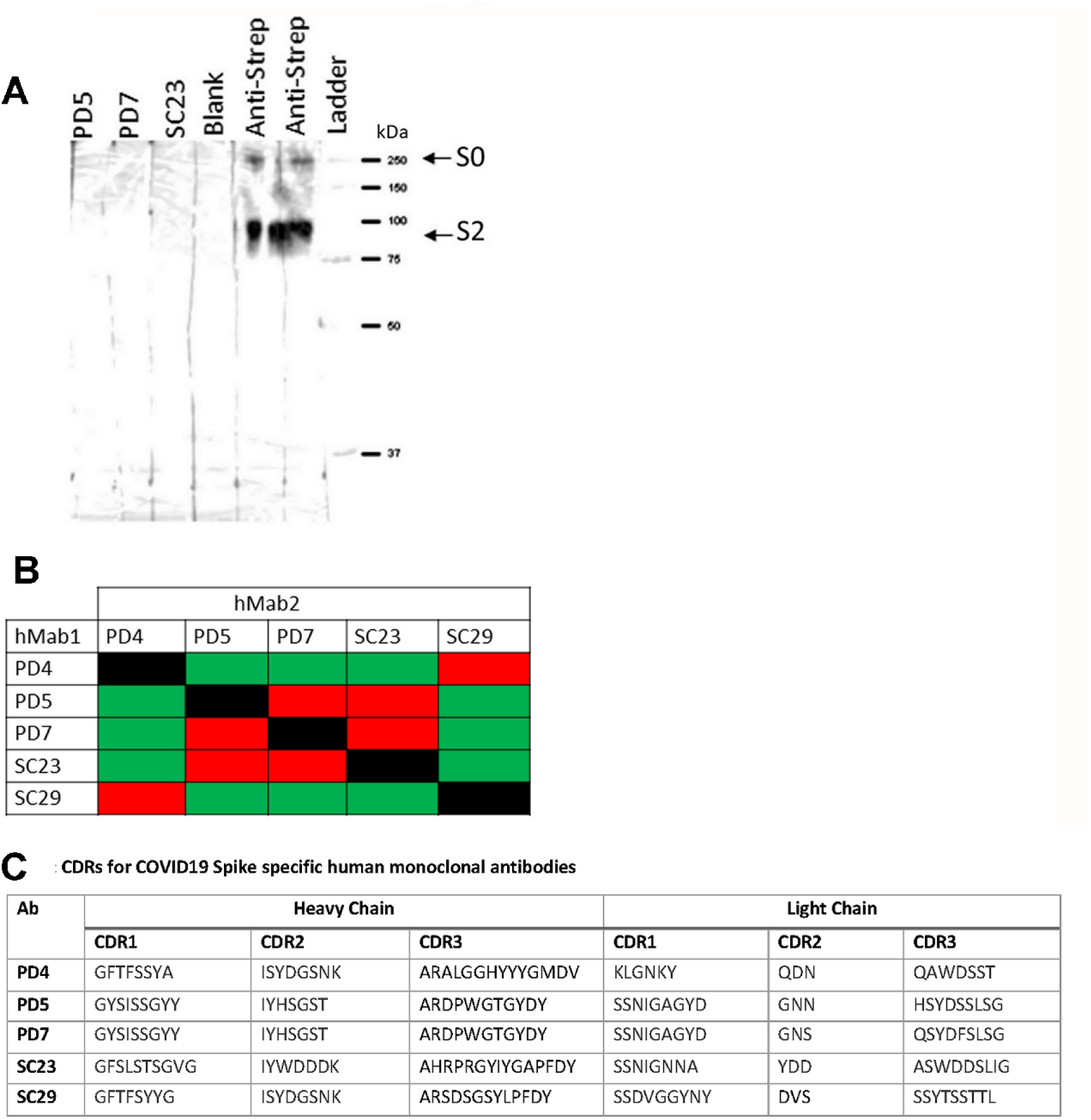
The binding activity of the human monoclonal antibodies on the SARS-CoV-2 S protein. **(A)** The recombinant SARS-CoV-2 S-Strep protein was transferred on to a polyvinyl difluoride (PVDF) membrane by Western blotting and the PVDF membrane was cut into strips and incubated with PD5, PD7 or SC23, followed by secondary antibodies conjugated with horse radish peroxidase (HRP). The lane representing blank refers to a blank lane and the lanes representing Anti-Strep refers to incubation with Anti-Strep conjugated to HRP as is the loading control (positive S protein control). The protein species corresponding in size to the uncleaved S0 and S2 are highlighted. The Ladder represents protein standards from 37 to 250 kDa. **(B)** In the ELISA-based antibody binding competition assay each of the hMAbs (PD4, PD5, PD7, SC23, SC29) was coated in a well (hMAb1) and the recombinant S protein bound to the coated antibody. Different hMAbs (hMab2) were then added to the bound S protein and binding assessed by ELISA. Red = binding of the specific hMab1 inhibits the specific hMab2 antibody binding; while green = binding of the specific hMab1 has no effect on the specific hMab2 antibody binding. **(C)** The sequences of the complementarity-determining regions (CDRs) for member of the antibody panel. The CDR1-3 sequences are shown for the heavy and light chain for each corresponding antibody.

**SFig. 3.**
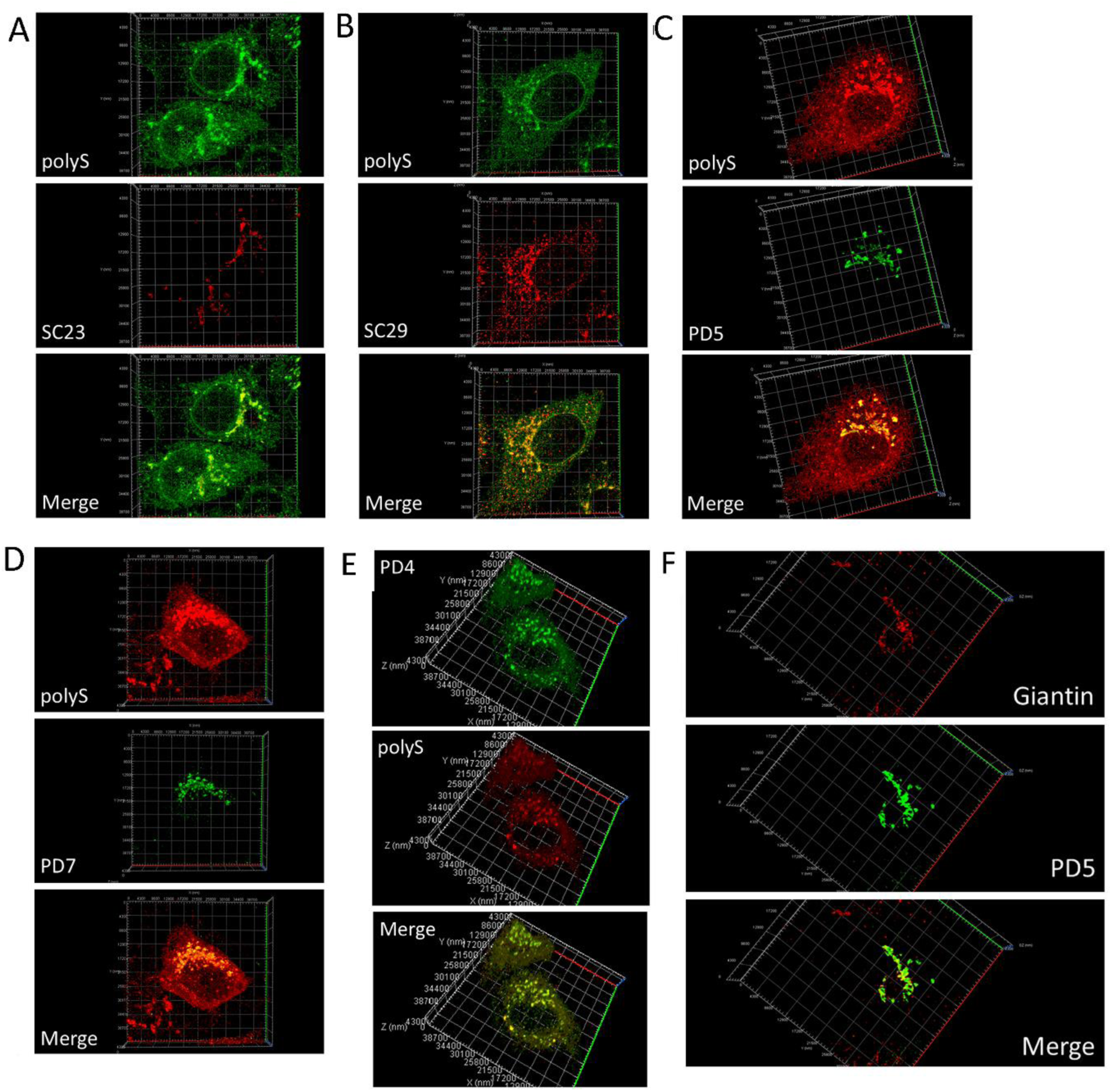
The reconstructed Z-stack images obtained by confocal microscopy from virus-infected Vero E6 cells stained with the anti-SARS-CoV-2 human monoclonal antibodies. At 18 hrs post-infection (hpi) SARS-CoV-2/0334-infected Vero E6 cells were co-stained with anti-polyS and either **(A)** SC23, **(B)** SC29, **(C)** PD5, **(D)** PD7 and **(E)** PD4, and imaged using confocal microscopy. A series of images from the same cell were obtained in the Z-projection and the individual images reconstructed into 3-dimentional images of the co-stained cells. **(F)** At 18 hpi SARS-CoV-2/0334-infected cells were co-stained with PD5 and anti-giantin, and imaged using confocal microscopy. A series of images from the same cell were obtained in the Z-projection and the individual images reconstructed into 3-dimentional images of the co-stained cells.

**SFig. 4.**
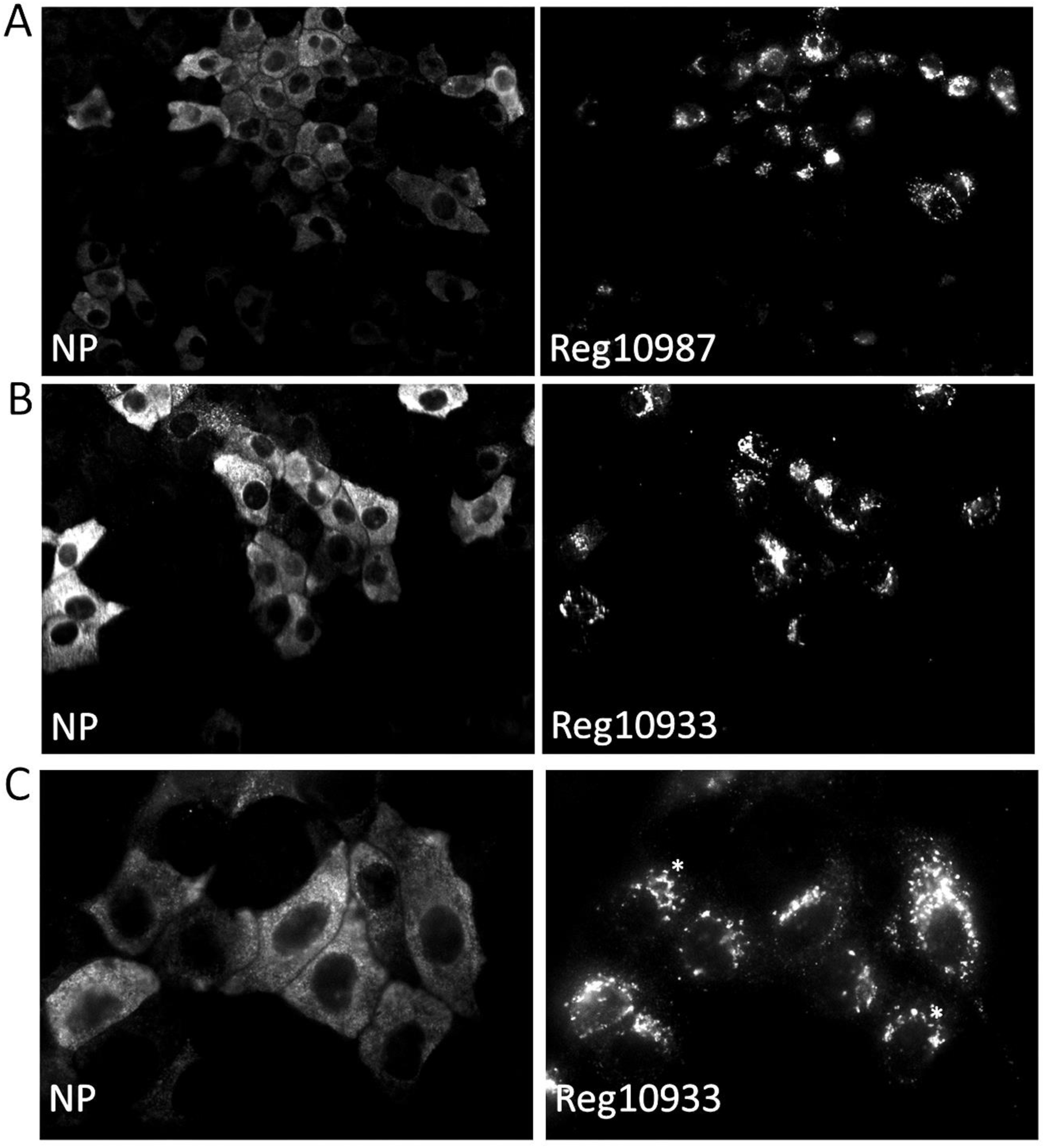
Vero E6 cells infected with SARS-CoV-2 and stained with monoclonal antibodies with whose RDB recognition sites have been defined. SARS-CoV-2/0334-infected Vero E6 cells were co-stained with either **(A)** REGEN-10987 (Reg10987) or **(B and C)** REGEN-10933 (Reg10933) and anti-NP (recognizes the SARS-CoV-2 N protein). In all cases images were recorded using immunofluorescence microscopy. Plates (A) and (B); (objective x20 magnification) and plate (C) (objective x100 magnification).

**SFig. 5.**
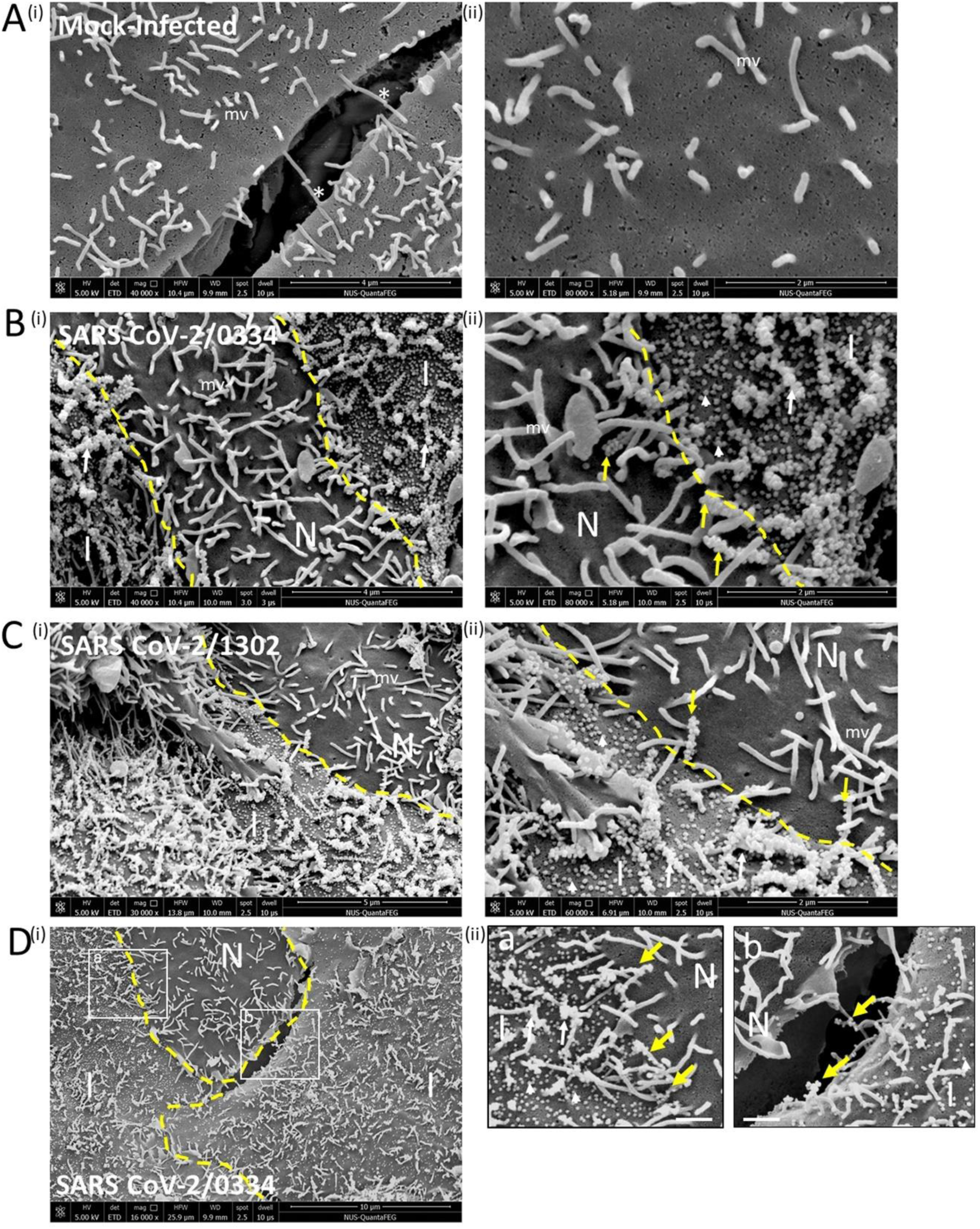
Direct evidence for cell-to-cell transmission of SARS-CoV-2 particles via cell surface projections in virus-infected Vero E6 cells. **(A)** Mock-infected Vero E6 cells and cells infected with **(B)** SARS-CoV-2/0334, **(C)** SARS-CoV-2/1302 and **(D)** SARS-CoV-2/0334 were imaged using scanning electron microscopy at 18 hrs post-infection. In each case microvilli (mv), infected (I) and non-infected (N) cells, the borders between different cells (broken yellow line), individual virus particles (white arrow head), virus particles on microvilli (white arrows), virus particles on microvilli spanning different cells (yellow arrow) and microvilli spanning different cells in mock-infected cell monolayer (*) are highlighted with **A**(i) and **B**(i) at 40,000x magnification, **A**(ii), **B**(ii) and **C**(ii) at 80,000x magnification, **C**(i) at 30,000x magnification, **C**(ii) at 60,000x magnification, and **D**(i) at 16,000x magnification. **D**(ii) Plates a and b are enlarged images from the area demarcated by the open white boxes in **D**(i). The white bars in **D**(ii) represent 1µm.

**SFig. 6.**
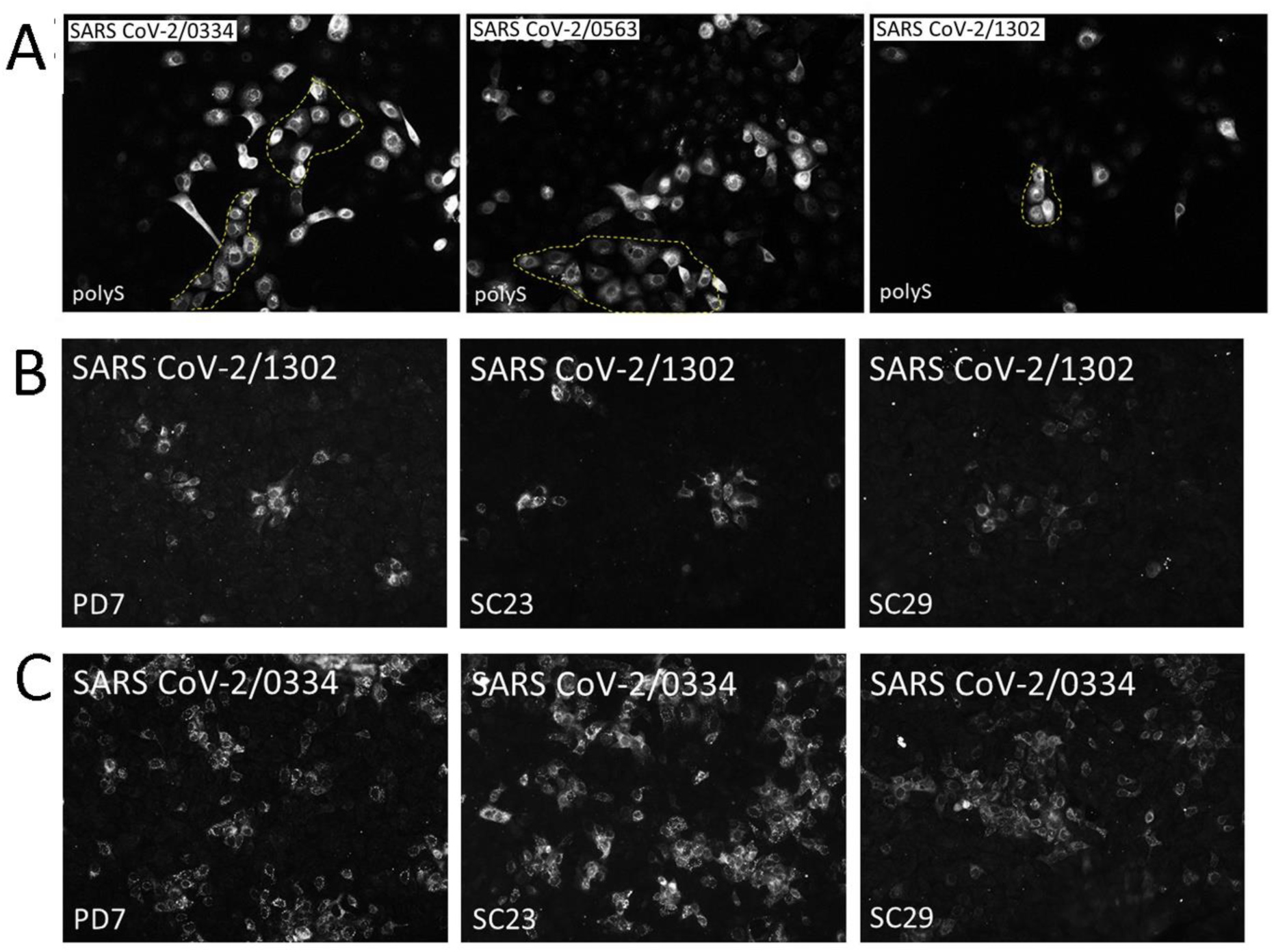
Vero E6 cells infected with SARS-CoV-2 at low multiplicity of infection and stained with PD7, SC23, SC29 and polyS. **(A)** Cells were infected with SARS-CoV-2/0334, SARS-CoV-2/0563 and SARS-CoV-2/1302 as indicated using a multiplicity of infection (moi) of 0.01. At 18 hrs post-infection the cells were stained with polyS. In all cases images were recorded using immunofluorescence microscopy (objective x20 magnification). Cells were infected with **(B)** SARS-CoV-2/1302 and (C) SARS-CoV-2/0334 using a multiplicity of infection (moi) of 0.01 and at 18 hpi the cells were stained with PD7, SC23 and SC29 as indicated.

**SFig. 7.**
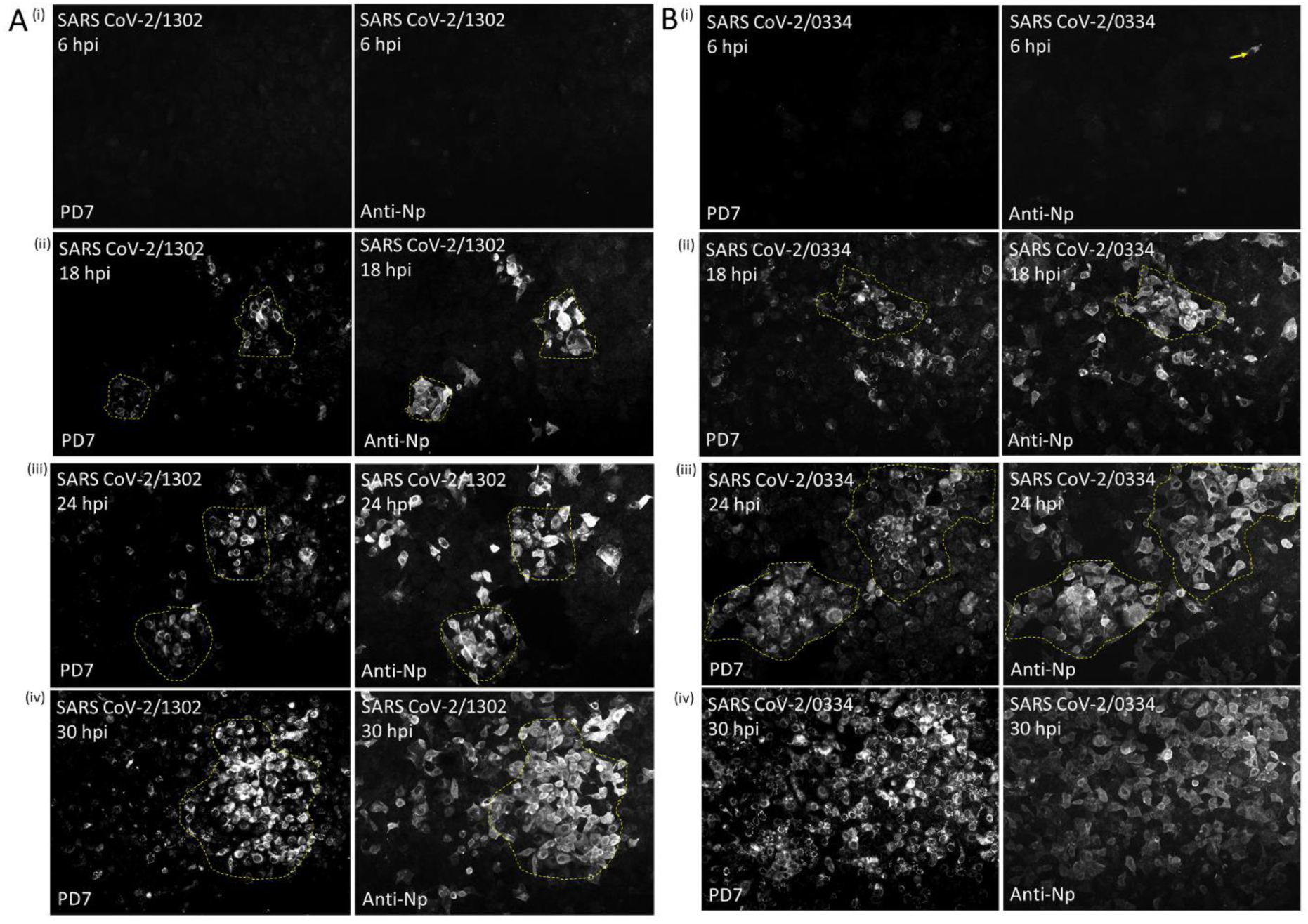
Temporal appearance of PD7 and anti-N protein staining on Vero E6 cells infected with SARS-CoV-2/1302 and SARS-CoV-2/0334. Cells were infected with **(A)** SARS-CoV-2/1302 and **(B)** SARS-CoV-2/0334 using a multiplicity of infection of 0.01 and at (i) 6 hrs post-infection (hpi), (ii) 18 hpi, (iii) 24 hpi and (iv) 30 hpi the cells were co-stained with PD7 and anti-N protein (NP). The stained cells were imaged using immunofluorescence microscopy (objective x20). In both **(A)** and **(B),** clusters of infected cells are highlighted (broken yellow line).

**SFig. 8.**
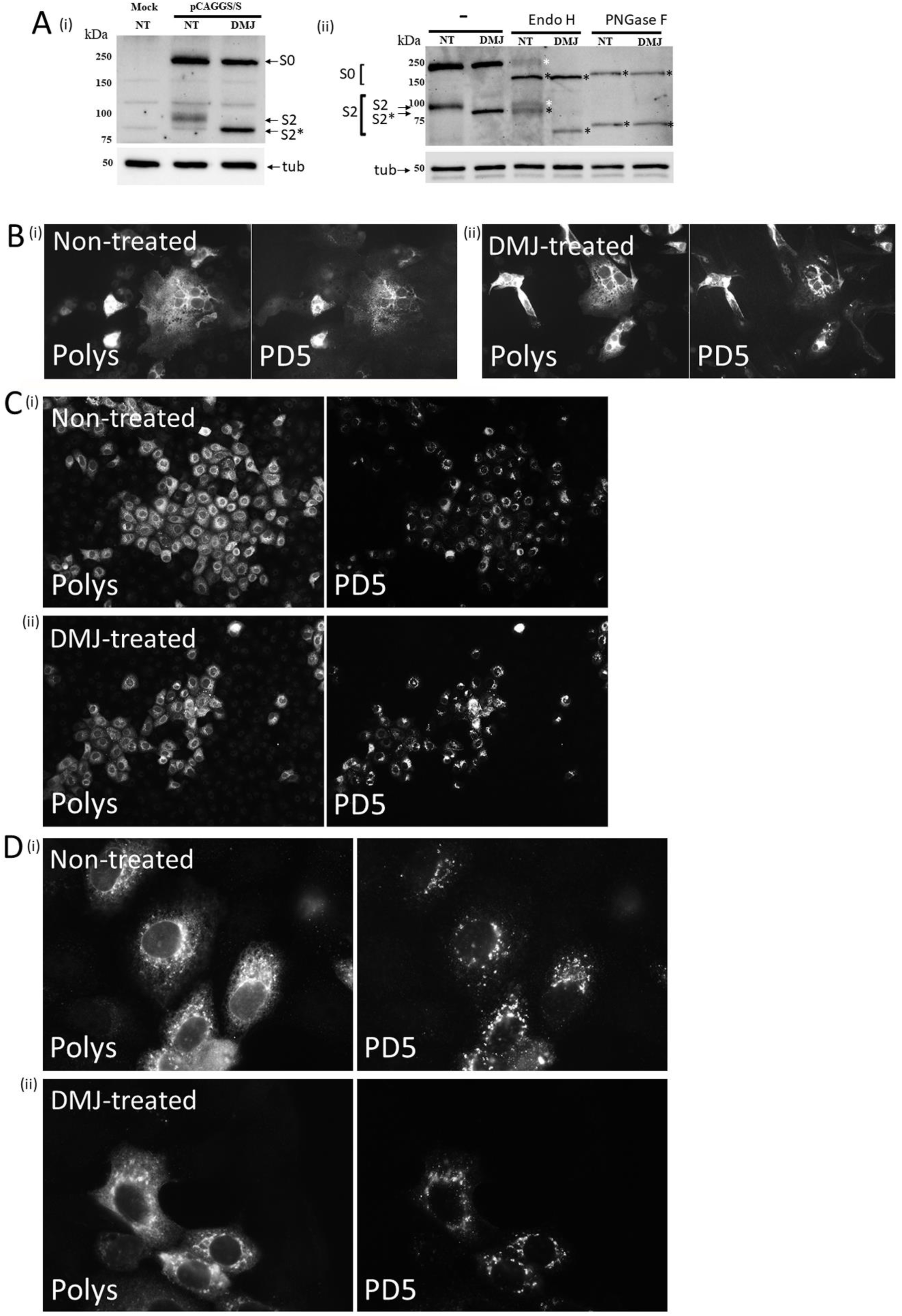
Maturation of the N-linked glycans in the S protein RBD is not required for PD5 recognition. **(A) (i)** Cells were mock-transfected and transfected with pCAGGS/S in the absence (NT) and presence of 1mM deoxymannojirmycin (DMJ) and the cell lysates immunoblotted with polyS. Protein species corresponding in size to the uncleaved (S0) and S2 domain of the S protein are indicated. Also indicated (S2*) is the faster migrating S2 domain that is formed in the presence of DMJ. (ii) Cells expressing the S protein in non-treated (NT) and DMJ-treated cells were either mock-digested (Mock) or treated with EndoH and PNGase F and the samples immunoblotted with polyS. The migration of the EndoH resistant (white *) and EndoH and PNGaseF sensitive (black*) S protein species are indicated. In all cases immunoblotting with anti-tubulin(tub) is the loading control. **(B)** Non-treated (NT) and DMJ-treated cells expressing the recombinant S protein were co-stained with polyS and PD5 and imaged by IF microscopy (objective x40 magnification). At 18 hrs post-infection non-treated (NT) and DMJ-treated cells infected with SARS-CoV-2/0334 were co-stained with polyS and PD5 and imaged by IF microscopy with **(C)** objective x40 magnification and **(D)** objective x100 magnification.

**SFig 9.**
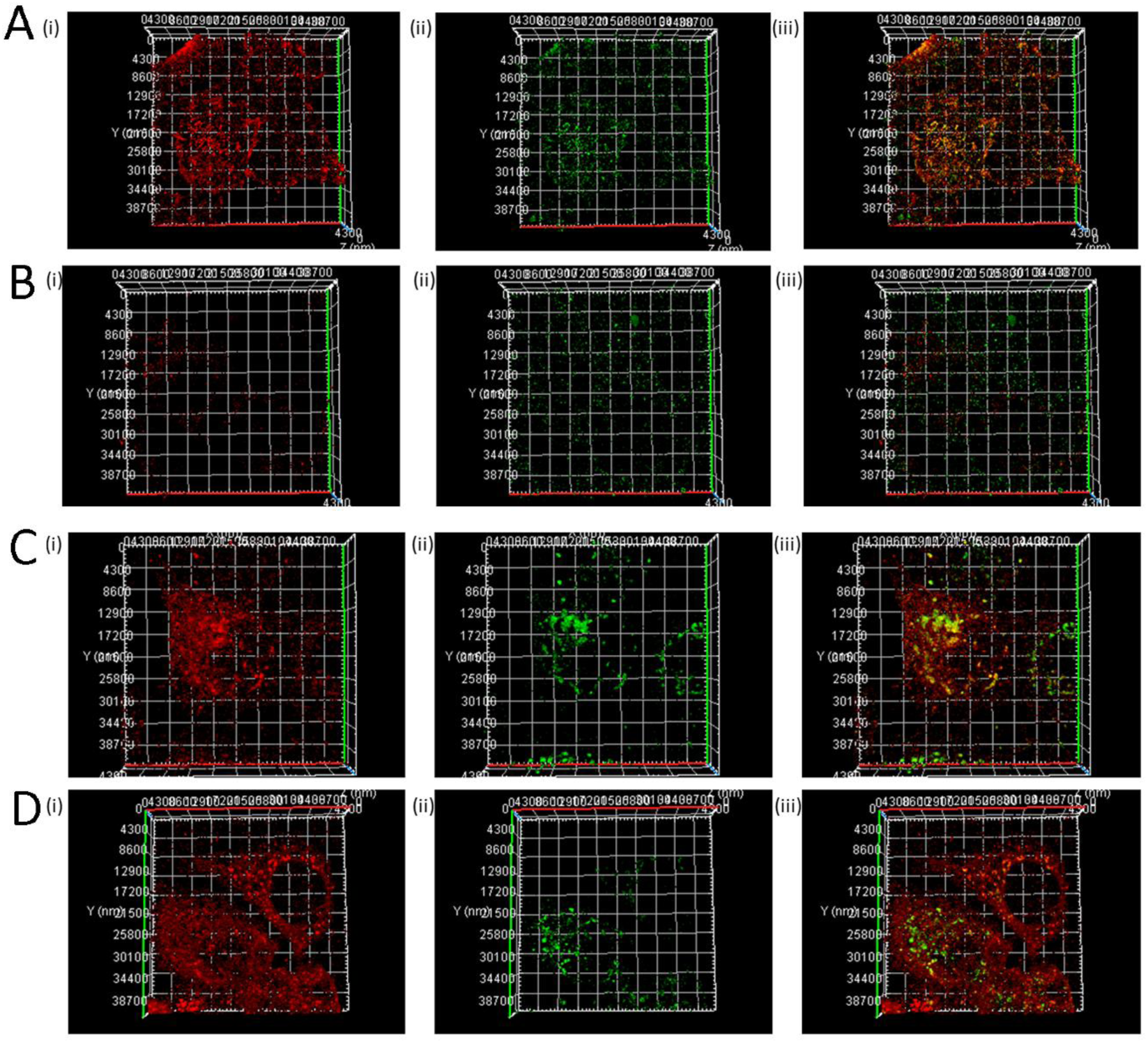
The reconstructed Z-stack images obtained by confocal microscopy from non-treated and decanoyl-RVKR-cmk-treated virus-infected Vero E6 cells stained with PD5. At 18 hrs post-infection, (i) non-treated and (ii) decanoyl-RVKR-cmk-treated SARS-CoV-2/0334-infected Vero E6 cells were **(A and B)** non-permeabilized and **(C and D)** permeabilized, and co-stained with polyS and PD5, and imaged using confocal microscopy. A series of images from the same cell were obtained in the Z-projection and the individual images reconstructed into 3-dimentional images of the co-stained cells.

## References

1. Li, Q., et al., Early Transmission Dynamics in Wuhan, China, of Novel Coronavirus-Infected Pneumonia. N Engl J Med, 2020. 382(13): p. 1199–1207.

2. Zhu, N., et al., A Novel Coronavirus from Patients with Pneumonia in China, 2019. N Engl J Med, 2020. 382(8): p. 727–733.

3. van der Hoek, L., Human coronaviruses: what do they cause? Antivir Ther, 2007. 12(4 Pt B): p. 651–8.

4. Drosten, C., et al., Identification of a novel coronavirus in patients with severe acute respiratory syndrome. N Engl J Med, 2003. 348(20): p. 1967–76.

5. Nickbakhsh, S., et al., Epidemiology of Seasonal Coronaviruses: Establishing the Context for the Emergence of Coronavirus Disease 2019. J Infect Dis, 2020. 222(1): p. 17–25.

6. Fehr, A.R. and S. Perlman, Coronaviruses: an overview of their replication and pathogenesis. Methods Mol Biol, 2015. 1282: p. 1–23.

7. Zhou, P., et al., A pneumonia outbreak associated with a new coronavirus of probable bat origin. Nature, 2020. 579(7798): p. 270–273.

8. Veneti, L., et al., Increased risk of hospitalisation and intensive care admission associated with reported cases of SARS-CoV-2 variants B.1.1.7 and B.1.351 in Norway, December 2020 - May 2021. PLoS One, 2021. 16(10): p. e0258513.

9. Bager, P., et al., Risk of hospitalisation associated with infection with SARS-CoV-2 lineage B.1.1.7 in Denmark: an observational cohort study. Lancet Infect Dis, 2021. 21(11): p. 1507–1517.

10. Challen, R., et al., Risk of mortality in patients infected with SARS-CoV-2 variant of concern 202012/1: matched cohort study. Bmj, 2021. 372: p. n579.

11. Hu, B., et al., Characteristics of SARS-CoV-2 and COVID-19. Nat Rev Microbiol, 2021. 19(3): p. 141–154.

12. Tang, T., et al., Proteolytic Activation of SARS-CoV-2 Spike at the S1/S2 Boundary: Potential Role of Proteases beyond Furin. ACS Infect Dis, 2021. 7(2): p. 264–272.

13. Bertram, S., et al., TMPRSS2 and TMPRSS4 facilitate trypsin-independent spread of influenza virus in Caco-2 cells. J Virol, 2010. 84(19): p. 10016–25.

14. Simmons, G., et al., Inhibitors of cathepsin L prevent severe acute respiratory syndrome coronavirus entry. Proc Natl Acad Sci U S A, 2005. 102(33): p. 11876–81.

15. Kabir, M.A., et al., Management of COVID-19: current status and future prospects. Microbes Infect, 2021. 23(4-5): p. 104832.

16. Brouwer, P.J.M., et al., Potent neutralizing antibodies from COVID-19 patients define multiple targets of vulnerability. Science, 2020. 369(6504): p. 643–650.

17. Jennewein, M.F., et al., Isolation and Characterization of Cross-Neutralizing Coronavirus Antibodies from COVID-19+ Subjects. bioRxiv, 2021.

18. Zost, S.J., et al., Potently neutralizing and protective human antibodies against SARS-CoV-2. Nature, 2020. 584(7821): p. 443–449.

19. Chan, C.E.Z., et al., The Fc-mediated effector functions of a potent SARS-CoV-2 neutralizing antibody, SC31, isolated from an early convalescent COVID-19 patient, are essential for the optimal therapeutic efficacy of the antibody. PLoS One, 2021. 16(6): p. e0253487.

20. Baum, A., et al., REGN-COV2 antibodies prevent and treat SARS-CoV-2 infection in rhesus macaques and hamsters. Science, 2020. 370(6520): p. 1110–1115.

21. Weinreich, D.M., et al., REGN-COV2, a Neutralizing Antibody Cocktail, in Outpatients with Covid-19. N Engl J Med, 2021. 384(3): p. 238–251.

22. Wrobel, A.G., et al., SARS-CoV-2 and bat RaTG13 spike glycoprotein structures inform on virus evolution and furin-cleavage effects. Nat Struct Mol Biol, 2020. 27(8): p. 763–767.

23. Wrapp, D., et al., Cryo-EM structure of the 2019-nCoV spike in the prefusion conformation. Science, 2020. 367(6483): p. 1260–1263.

24. Chia, P.Y., et al., Detection of air and surface contamination by SARS-CoV-2 in hospital rooms of infected patients. Nat Commun, 2020. 11(1): p. 2800.

25. Notredame, C., D.G. Higgins, and J. Heringa, T-Coffee: A novel method for fast and accurate multiple sequence alignment. J Mol Biol, 2000. 302(1): p. 205–17.

26. Gasteiger, E., et al., ExPASy: The proteomics server for in-depth protein knowledge and analysis. Nucleic Acids Res, 2003. 31(13): p. 3784–8.

27. de Haard, H.J., et al., A large non-immunized human Fab fragment phage library that permits rapid isolation and kinetic analysis of high affinity antibodies. J Biol Chem, 1999. 274(26): p. 18218–30.

28. Jeffree, C.E., et al., Distribution of the attachment (G) glycoprotein and GM1 within the envelope of mature respiratory syncytial virus filaments revealed using field emission scanning electron microscopy. Virology, 2003. 306(2): p. 254–67.

29. Andrés, C., et al., Naturally occurring SARS-CoV-2 gene deletions close to the spike S1/S2 cleavage site in the viral quasispecies of COVID19 patients. Emerg Microbes Infect, 2020. 9(1): p. 1900–1911.

30. Lontok, E., E. Corse, and C.E. Machamer, Intracellular targeting signals contribute to localization of coronavirus spike proteins near the virus assembly site. J Virol, 2004. 78(11): p. 5913–22.

31. Copin, R., et al., The monoclonal antibody combination REGEN-COV protects against SARS-CoV-2 mutational escape in preclinical and human studies. Cell, 2021. 184(15): p. 3949–3961.e11.

32. Starr, T.N., et al., Prospective mapping of viral mutations that escape antibodies used to treat COVID-19. Science, 2021. 371(6531): p. 850–854.

33. Zhao, P., et al., Virus-Receptor Interactions of Glycosylated SARS-CoV-2 Spike and Human ACE2 Receptor. Cell Host Microbe, 2020. 28(4): p. 586–601.e6.

34. Laue, M., et al., Morphometry of SARS-CoV and SARS-CoV-2 particles in ultrathin plastic sections of infected Vero cell cultures. Sci Rep, 2021. 11(1): p. 3515.

35. Zeng, C., et al., SARS-CoV-2 Spreads through Cell-to-Cell Transmission. bioRxiv, 2021.

36. Caldas, L.A., et al., Ultrastructural analysis of SARS-CoV-2 interactions with the host cell via high resolution scanning electron microscopy. Sci Rep, 2020. 10(1): p. 16099.

37. Hirano, N., K. Fujiwara, and M. Matumoto, Mouse hepatitis virus (MHV-2). Plaque assay and propagation in mouse cell line DBT cells. Jpn J Microbiol, 1976. 20(3): p. 219–25.

38. Harcourt, J., et al., Isolation and characterization of SARS-CoV-2 from the first US COVID-19 patient. bioRxiv, 2020.

39. Cai, Y., et al., Distinct conformational states of SARS-CoV-2 spike protein. Science, 2020. 369(6511): p. 1586–1592.

40. Lozada, C., et al., Identification and Characteristics of Fusion Peptides Derived From Enveloped Viruses. Front Chem, 2021. 9: p. 689006.

41. Molloy, S.S., et al., Intracellular trafficking and activation of the furin proprotein convertase: localization to the TGN and recycling from the cell surface. Embo j, 1994. 13(1): p. 18–33.

42. McDonald, T.P., et al., Evidence that maturation of the N-linked glycans of the respiratory syncytial virus (RSV) glycoproteins is required for virus-mediated cell fusion: The effect of alpha-mannosidase inhibitors on RSV infectivity. Virology, 2006. 350(2): p. 289–301.

43. McDonald, T.P. and R.J. Sugrue, The use of two-dimensional SDS-PAGE to analyze the glycan heterogeneity of the respiratory syncytial virus fusion protein. Methods Mol Biol, 2007. 379: p. 97–108.

44. Miller, L.M., et al., Heterogeneity of Glycan Processing on Trimeric SARS-CoV-2 Spike Protein Revealed by Charge Detection Mass Spectrometry. J Am Chem Soc, 2021. 143(10): p. 3959–3966.

45. Watanabe, Y., et al., Site-specific glycan analysis of the SARS-CoV-2 spike. Science, 2020. 369(6501): p. 330–333.

46. Yao, H., et al., Molecular Architecture of the SARS-CoV-2 Virus. Cell, 2020. 183(3): p. 730–738.e13.

47. Sugrue, R.J., et al., Furin cleavage of the respiratory syncytial virus fusion protein is not a requirement for its transport to the surface of virus-infected cells. J Gen Virol, 2001. 82(Pt 6): p. 1375–86.

48. Papa, G., et al., Furin cleavage of SARS-CoV-2 Spike promotes but is not essential for infection and cell-cell fusion. PLoS Pathog, 2021. 17(1): p. e1009246.

49. Xia, S., et al., The role of furin cleavage site in SARS-CoV-2 spike protein-mediated membrane fusion in the presence or absence of trypsin. Signal Transduct Target Ther, 2020. 5(1): p. 92.

50. Gao, S., et al., ACE2 isoform diversity predicts the host susceptibility of SARS-CoV-2. Transbound Emerg Dis, 2021. 68(3): p. 1026–1032.

51. Conceicao, C., et al., The SARS-CoV-2 Spike protein has a broad tropism for mammalian ACE2 proteins. PLoS Biol, 2020. 18(12): p. e3001016.

52. Damas, J., et al., Broad host range of SARS-CoV-2 predicted by comparative and structural analysis of ACE2 in vertebrates. Proc Natl Acad Sci U S A, 2020. 117(36): p. 22311–22322.

53. Peacock, T.P., et al., The furin cleavage site in the SARS-CoV-2 spike protein is required for transmission in ferrets. Nat Microbiol, 2021. 6(7): p. 899–909.

54. Johnson, B.A., et al., Loss of furin cleavage site attenuates SARS-CoV-2 pathogenesis. Nature, 2021. 591(7849): p. 293–299.

55. Duan, L., et al., The SARS-CoV-2 Spike Glycoprotein Biosynthesis, Structure, Function, and Antigenicity: Implications for the Design of Spike-Based Vaccine Immunogens. Front Immunol, 2020. 11: p. 576622.

56. Harvey, W.T., et al., SARS-CoV-2 variants, spike mutations and immune escape. Nat Rev Microbiol, 2021. 19(7): p. 409–424.

57. Bangaru, S., et al., Structural analysis of full-length SARS-CoV-2 spike protein from an advanced vaccine candidate. Science, 2020. 370(6520): p. 1089–1094.

58. Turoňová, B., et al., In situ structural analysis of SARS-CoV-2 spike reveals flexibility mediated by three hinges. Science, 2020. 370(6513): p. 203–208.

